# Fingerprint SRS Imaging Unveils Ergosteryl Ester as a Metabolic Signature of Azole-Resistant *Candida albicans*

**DOI:** 10.1101/2022.12.09.519679

**Authors:** Meng Zhang, Pu-Ting Dong, Hassan E. Eldesouky, Yuewei Zhan, Haonan Lin, Zian Wang, Ehab A. Salama, Sebastian Jusurf, Cheng Zong, Zhicong Chen, Mohamed N. Seleem, Ji-Xin Cheng

## Abstract

*Candida albicans* (*C. albicans*), a major fungal pathogen, causes life-threatening infections in immunocompromised individuals. Fluconazole (FLC) is recommended as first-line therapy for treatment of invasive fungal infections. Yet, the widespread use of FLC has resulted in increased antifungal resistance among different strains of *Candida*, especially *C. albicans*, which is a leading source of hospital-acquired infections. Here, by hyperspectral stimulated Raman scattering (hSRS) imaging of single fungal cells in the fingerprint window and pixel-wise spectral unmixing, we report aberrant ergosteryl ester accumulation in azole-resistant *C. albicans* compared to azole-susceptible species. This accumulation was a consequence of *de novo* lipogenesis. Lipid profiling by mass spectroscopy identified ergosterol oleate to be the major species stored in azole-resistant *C. albicans*. Blocking ergosterol esterification by oleate and suppressing sterol synthesis by FLC synergistically suppressed the viability of *C. albicans in vitro* and limited the growth of biofilm on mouse skin *in vivo*. Our findings highlight a metabolic marker and a new therapeutic strategy for targeting azole-resistant *C. albicans* by interrupting the esterified ergosterol biosynthetic pathway.

**Significance Statement:** Invasive fungal infections and increasing antifungal resistance are emerging threats to public health with high morbidity and mortality. Despite the advances in azole resistance mechanisms, it remains unclear why some fungal species are intrinsically resistant to or easily acquire resistance to multiple antifungal drugs. Here, using fingerprint SRS microscopy, we uncovered a molecular signature, aberrant ergosteryl ester accumulation, linked to the azole resistance of *Candida* species. An antifungal treatment strategy combining oleate (inhibitor of ersgosteryl esterification) and azole significantly attenuates the azole resistance and the viability of *C. albicans in vitro* and *in vivo*. Our work opens a new way to detect and treat azole-resistant fungal infections by targeting ergosterol metabolism.

## Introduction

Invasive fungal infections and increasing resistance to antifungals are emerging threats to public health that contribute to high morbidity and mortality (1). Fungal infections have been referred to as “hidden killers,” because the effects of fungal infections and antifungal resistance on human health are not widely recognized (2). *Candida albicans* (*C. albicans*) is a major fungal pathogen that causes life-threatening infections when the host becomes debilitated or immunocompromised (3). Species of *Candida*, most notably *C. albicans*, are mostly associated with invasive, life-threatening fungal infections in immunocompromised individuals (4). Mortality rates due to fungal infections are estimated to be as high as 45% (5), which may be due to inefficient diagnostic methods and inappropriate initial antifungal therapies (6).

Therapeutic options for fungal infections are limited. The most widely used antifungal drugs comprise only a few chemical classes, including azoles (fluconazole, itraconazole, voriconazole, and posaconazole), polyenes (amphotericin B), and the echinocandins (caspofungin, anidulafungin, and micafungin) (7, 8). Azoles are recommended as first-line therapy for most invasive *Candida* species that cause systemic infections; azoles inhibit 14α-demethylase *Erg11* in the ergosterol biosynthesis pathway. This results in the accumulation of toxic sterol 14,24-dimethylcholesta-8,24(28)-dien-3β,6α-diol (DMCDD), which permeabilizes the fungal plasma membrane (9). However, the widespread use of azoles has resulted in increased antifungal resistance by different fungal strains to these drugs, especially among *Candida* species (10, 11). *C. albicans* can gain resistance to azoles mainly via genetic alteration of the drug target *Erg11* (12); upregulation of the efflux pumps *CDR1, CDR2*, and *MDR1* (13-15); and inactivation of *ERG3*, which synthesizes the sterol (10, 11, 16-21). Despite these advances in our understanding of azoles resistance mechanisms, it remains unclear why some fungal species are intrinsically resistant to or easily acquire resistance to multiple antifungal drugs (1, 22). In particular, how ergosterol metabolism is reprogrammed in response to antifungal azole treatment is poorly understood.

Recently developed coherent Raman scattering microscopy, based on coherent anti-Stokes Raman scattering (CARS) or stimulated Raman scattering (SRS), opens a new window to explore single cell metabolism in a spatially and temporally resolved manner. In particular, hyperspectral CARS or SRS imaging has unveiled hidden signatures in various biological systems. These imaging techniques have permitted researchers to spatially resolve and quantitatively analyze metabolites inside cancer cells (23-28) and *Caenorhabditis elegans* (29-34). Dynamic imaging of specific metabolites was enabled by SRS imaging of vibrational probes (35-38).

As it relates to using SRS imaging for infectious diseases, the orientation of amphotericin B was resolved by polarization-sensitive SRS signal from fingerprint C=C stretching vibration (39). Rapid antimicrobial susceptibility determination at a single-bacterium level was achieved by stimulated Raman metabolic imaging (40, 41). Despite these advances, SRS imaging of metabolism in drug-resistant fungal cells is underexplored. A recent femtosecond SRS study identified lipid accumulation in azole-resistant cells (42). Yet, femtosecond SRS in the CH stretching vibration window does not have the capacity to resolve the chemical content of lipids. Consequently, the molecular mechanism and clinical impact of this lipid accumulation remains elusive.

To study metabolic reprogramming of fungal cells in response to azole treatment, we employed fingerprint hyperspectral SRS (hSRS) imaging to visualize the contents of *C. albicans* at a subcellular level. A pixel-wise least absolute shrinkage and selection operator (LASSO) regression algorithm was further applied to decompose the hSRS stack into chemical maps. An aberrant storage of esterified ergosterol (EE), featured by the sterol C=C peak at 1603 cm^−1^ and the acyl C=C peak at 1655 cm^−1^, was identified in azole-resistant species, but not in azole-sensitive species. Further investigation verified that EE accumulation in azole-resistant *C. albicans* arises from *de novo* lipogenesis. Mass spectrometry analysis identified ergosteryl oleate as the major EE species. Based on these findings, we tested an antifungal strategy utilizing oleate to interrupt the esterification process. Oleate significantly suppressed EE accumulation in *C. albicans*. Moreover, oleate/azole combination treatment resulted in effective attenuation of the azole tolerance and viability of *C. albicans* in both yeast and biofilm forms. The *in vivo* study further confirmed that oleate-mediated inhibition of EE accumulation effectively impaired azole resistance in *C. albicans* and suppressed biofilm growth. These data collectively demonstrate the potential of using EE as a metabolic marker for detection of azole-resistant fungi and identify a new approach to treat invasive fungal infections by targeting ergosterol metabolism.

## Results

### SRS imaging reveals an increased level of esterified ergosteryl in azole-resistant *C. albicans*

We first applied confocal fluorescence imaging to confirm the accumulation of neutral lipids in stationary phase, fluconazole-resistant *C. albicans*. As shown in *SI Appendix*, Fig. S1, BDIPY-labeled droplets are seen in the *C. albicans* cells in the UPC (susceptible dose-dependent), TWO7241, and TWO7243 (resistant) strains, but are not seen in sensitive wild-type (W. Type) and DBC 46 strains. However, compositional information of individual lipid droplets (LDs) cannot be revealed from the fluorescence images. To quantitatively visualize and identify the chemical components of the lipids in individual fungal cells, we deployed fingerprint hSRS imaging via spectral focusing using a setup shown in *SI Appendix*, Fig. S2. SRS is a dissipative process in which energy corresponding to the beating frequency (ω_p_ − ω_S_) is transferred from input photons to a Raman-active molecular vibration (Ω). Tuning the time delay between the two chirped excitation beams can substantially change the overlapping difference in frequency, which excites different Raman shifts (Fig. 1A). By tuning the laser-beating frequency to cover the C=C stretching vibration window from 1550 to 1700 cm^−1^, we conducted hyperspectral SRS imaging of azole-resistant *C. albicans* strains, including TWO7241, TWO7243, NR-29446, ATCC 64124, ATCC MYA573, NR-29448, and azole-susceptible W. Type, all in stationary phase. The SRS spectra in this spectral region, which arise from the intracellular LDs and proteins, can be extracted at each pixel from the image stack. In the normalized SRS spectra of LDs in azole-resistant *C. albicans* (TWO7241, TWO7243, NR-29446, ATCC 64124), 2 strong Raman bands at 1603 cm^−1^ and 1655 cm^−1^ were present (Fig. 1B). The sterol C=C peak was absent in strain ATCC MYA573, which contains a mutation in *ERG11*. In comparison, the azole-sensitive *C. albicans* wild-type strain had a significantly weaker Raman signal at 1603 cm^−1^, which suggests that azole-susceptible *C. albicans* cells have a much lower concentration ratio of sterol C=C to acyl C=C (Fig. 1B). The 2 types of spectrally separated bands are contributed by the sterol C=C vibration with a peak at 1603 cm^−1^ and the acyl C=C vibration with a peak at 1655 cm^−1^, respectively. The origin of the 2 major peaks was confirmed by the SRS spectra of pure ergosterol and glyceryl trioleate, which exhibit a characteristic sterol C=C vibrational band at 1603 cm^−1^ and acyl C=C vibrational band at 1655 cm^−1^ (**Fig. 1C**). The SRS spectra of pure ergosterol and glyceryl trioleate overlapped with the spectra of LDs in azole-resistant *C. albicans* cells. This indicates that the content in individual LDs is predominantly in the form of ergosterol (in its esterified form) and glyceryl trioleate.

**Fig. 1.**
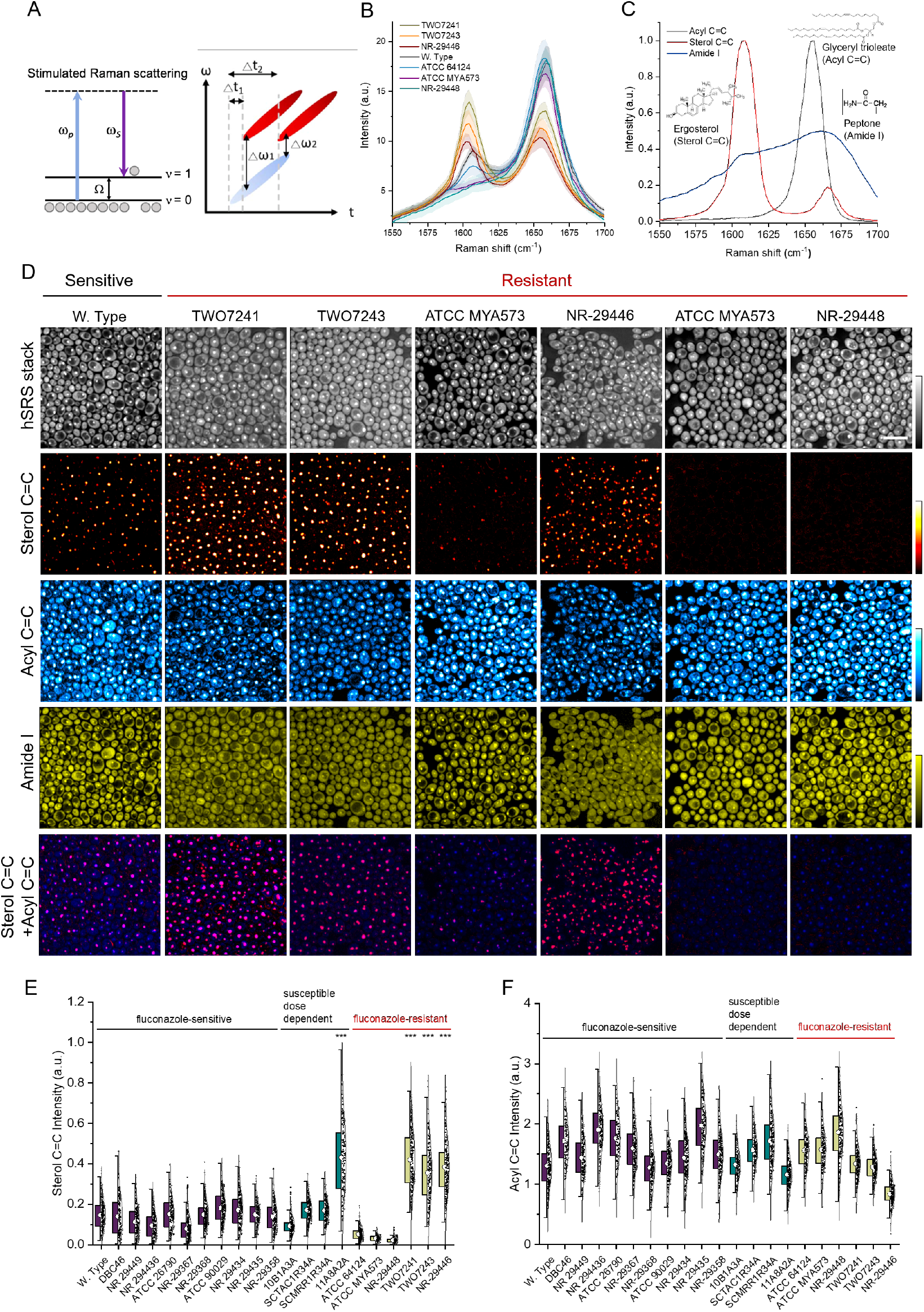
SRS imaging reveals an increased level of ergosteryl ester (EE) accumulation in azole-resistant *C. albicans*. (A) Schematic illustration on the principle of SRS. (B) Raman spectra of lipids accumulated in azole-resistant fungal strains, including *C. albicans* TWO7241, TWO7243, NR-29446, ATCC 64124, ATCC MYA573, NR-29448, and azole-susceptible *C. albicans* W. Type, by hyperspectral SRS imaging. (C) Reference spectra for hSRS spectra unmixing analysis using least square fitting. (D) Fingerprinting hyperspectral SRS images of various types of *C. albicans* cells, including azole-susceptible and azole-resistant cells. (E) EE and (F) Acyl C=C quantification analyses of (D). Bar scale represents 10 μm. Significance was evaluated using an unpaired *t*-test (***, p < 0.001).

To quantify the amount of EE in these LDs, concentration maps of acyl C=C, sterol C=C, and the amide I band were reconstructed from LASSO analysis of the hyperspectral SRS stacks (see methods). As shown in Fig. 1D, hyperspectral SRS images that contained hundreds of single fungal cells in each field-of-view were obtained. The standard reference spectra of ergosterol, glyceryl trioleate, and peptone (Fig. 1C) were used for unmixing of sterol C=C, acyl C=C, and the amide I band, respectively. The reconstructed concentration maps of sterol C=C, acyl C=C, amide I for azole-resistant *C. albicans* TWO7241, TWO7243, ATCC 64124, NR-29446, ATCC MYA573, NR-29448, and azole-susceptible *C. albicans* W. type are presented in Fig. 1D. The hSRS stack channel visualized the sum of hyperspectral SRS frames. Distinct spatial patterns were found in the decomposed sterol C=C, acyl C=C, and amide I channels. In the sterol C=C channel, EE accumulation was successfully separated and visualized in the *C. albicans* W. type, TWO7241, and TWO7243 strains, but barely in strains ATCC 64124, ATCC MYA573, and NR-29448. The acyl C=C signal revealed accumulation of lipid metabolites both in LDs and the cell membrane, whereas the amide I channel revealed protein distribution, which presented as a uniform pattern inside cells. The overlay image of the sterol C=C and acyl C=C concentration maps demonstrate the co-localization of the sterol C=C and acyl C=C LDs, which supports the fact that EE is stored with triglycerides in LDs.

To verify whether the observed phenomenon is strain specific, we repeated the detection on multiple azole-susceptible and susceptible dose-dependent *C. albicans* cells (*SI Appendix*, Fig. S3). Consistently, hyperspectral SRS spectral unmixing confirmed that azole-susceptible strains had significantly lower intracellular EE accumulation compared to azole-resistant strains. Interestingly, the UPC strain is a susceptible dose-dependent strain, but it exhibited obvious EE accumulation. It was found that the UPC strain, which is an *ERG11*-overexpressing isolates, contained a gain of function mutation in *UPC2*, in which 8 single amino acid substitutions were elucidated from their *UPC2* alleles. This was found to be associated with increased *ERG11* expression, increased ergosterol production, and decreased fluconazole susceptibility (11, 43).

For single cell chemical analysis, the decomposed concentration maps were segmented to generate maps of intracellular compartments corresponding to LDs and proteins in individual cells (*SI Appendix*, Fig. S4). Statistical analysis in Fig. 1E shows a clearly elevated level of sterol C=C accumulation in azole-resistant *C. albicans* TWO7241, TWO7243, and NR-29446. In contrast, the azole-resistant *C. albicans* strains ATCC 64124, ATCC MYA573, and NR-29448 had relatively lower levels of sterol C=C, probably due to involvement other azole resistance mechanisms that do not rely predominantly on ergosterol overproduction. Quantitative analysis of the EE-to-protein ratio intensity confirmed a significant difference in EE accumulation levels between azole-resistant and azole-susceptible or susceptible dose-dependent *C. albicans* (*SI Appendix*, Fig. S5A). For further statistical comparison, a Student’s *t*-test found that the 2 subpopulations were statistically different (*p* value < 0.001) in terms of the levels of EE in azole-resistant and azole-susceptible cells. In contrast, no significant alteration was present in the acyl C=C contents between azole-resistant and azole-susceptible strains as shown in the quantitative analysis of acyl C=C intensity (Fig. 1F) and acyl C=C-to-protein ratio intensity (*SI Appendix*, Fig. S5B). This indicates that acyl C=C is not a molecular marker inside *C. albicans*. These data collectively demonstrate a significantly increased level of EE accumulation in azole-resistant *C. albicans* compared to non-resistant strains.

Next, to examine the effects of growth period on EE accumulation, we explored the phase-dependent changes in lipid metabolism during fungal growth. In stationary phase, yeast cells have a balanced rate of microbial death and new cell generation. The metabolic activities of stationary-phase cells are at equilibrium. However, logarithmic-phase yeast cells grow and divide rapidly with minimal reproductive time. In logarithmic-phase yeast cells, metabolism is the most active at this stage of a cell’s lifespan and, as a consequence, these cells are more sensitive to changes in their environment (44, 45). Yeast cells accumulate more lipids during the stationary phase (46). Fig. 1 demonstrates the increased level of EE accumulation in stationary-phase, azole-resistant *C. albicans*. To explore whether EE is accumulated in the log phase as well, azole-resistant *C. albicans* TWO7241 cells were grown and then harvested in mid-logarithmic phase and stationary phase, respectively. The hyperspectral SRS concentration maps suggest that the level of EE is significantly decreased in logarithmic-phase cells compared to stationary-phase cells (Fig. 2A and 2C). The intracellular sterol C=C and acyl C=C intensities in individual cells were distinctly higher in stationary-phase cells, as shown in the SRS spectra of the lipids (Fig. 2B and 2D). The integrated sterol C=C and acyl C=C intensity in individual cells was quantitatively analyzed and plotted as histograms (Fig. 2G and 2H). The results indicated that the sterol C=C and acyl C=C contents were higher in azole-resistant *C. albicans* at a single-cell level. Additionally, we collected stationary-phase *C. albicans* cells and then cultured them in fresh nutrient medium for 3 h. The hyperspectral SRS spectra showed decreased EE accumulation in the *C. albicans* cells after the medium was refreshed (*SI Appendix*, Fig. S6). The growth of microorganisms depends on the availability of nutrients in the surrounding medium. A previous study found that when the culture medium of stationary-phase *C. albicans* cells was switched, this induced rapid hydrolyzation of sterol esters to free sterol and fatty acids that were utilized for the biogenesis of membranes (46). These data collectively suggest that higher levels of EE accumulation are a distinct metabolic feature of azole-resistant *C. albicans* cells that are in stationary phase.

**Fig. 2.**
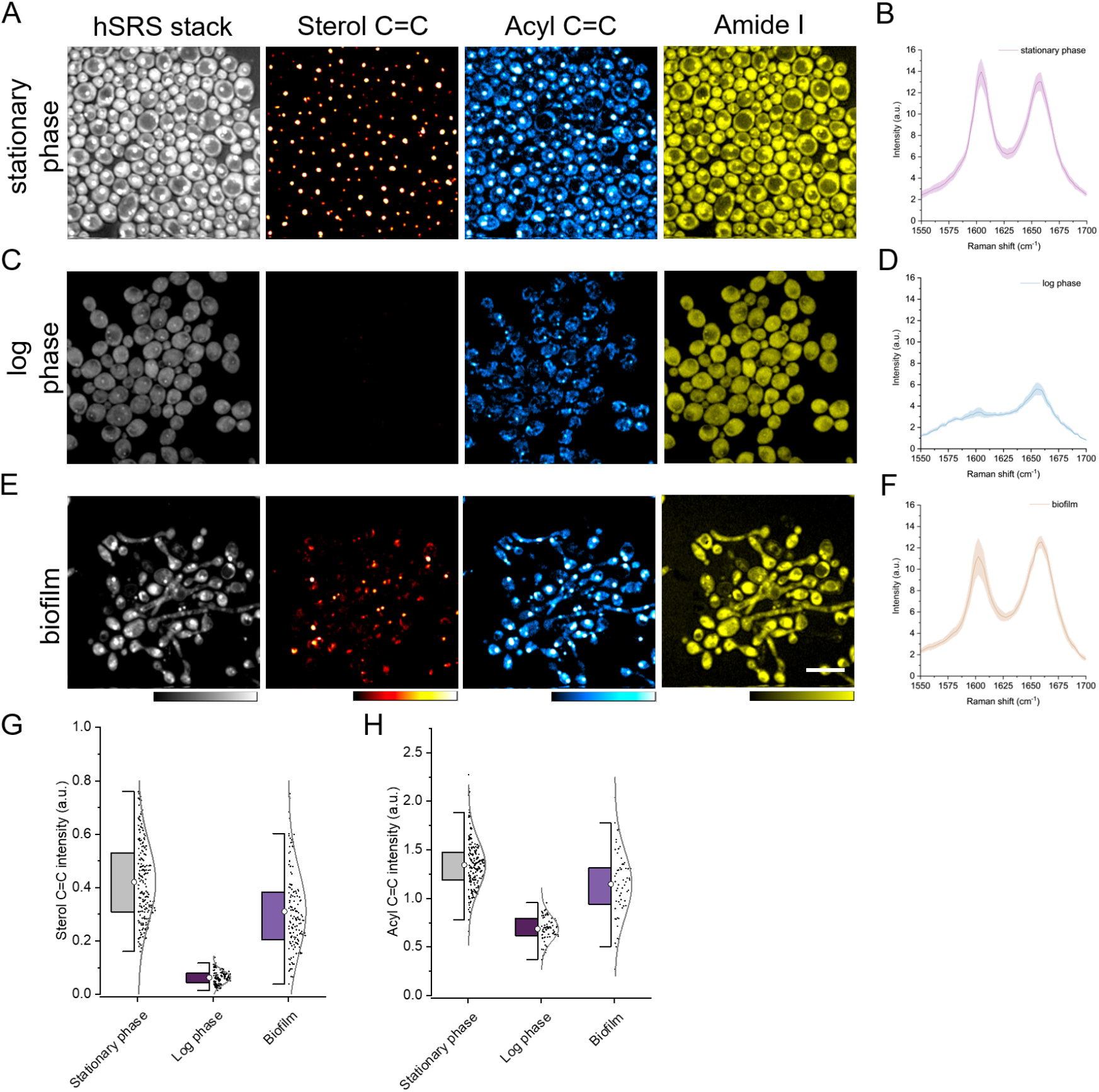
Increased EE accumulation in azole-resistant *C. albicans* stationary-phase and biofilm cells. Fingerprinting hyperspectral SRS images of stationary phase (A), logarithmic (log) phase (C) *C. albicans* cells, and *C. albicans* biofilm (E). hSRS spectra of lipids accumulated in (B) stationary phase, (D) logarithmic phase *C. albicans* TWO7241, and (F) *C. albicans* TWO7241 biofilm. (G) EE and (H) acyl C=C quantification analysis of hSRS unmixed concentration maps in (A), (C), and (E). Bar scale represents 10 μm.

*C. albicans* cells that are in stationary phase are capable of forming highly drug-resistant biofilms in humans through various adaptive mechanisms that alter the lipid composition of cell membranes. The ability of *C. albicans* to form biofilms poses a significant medical challenge in the treatment of candidiasis as these structured communities are recalcitrant to treatment by antifungals (47, 48). Therefore, we investigated if EE content is altered during *C. albicans* biofilm development. We cultured stationary-phase *C. albicans* to form biofilm and then examined the level of EE in cells using hSRS microscopy. A mixed type of cells, which comprised round and spherical yeast cells with filamentous hyphae and pseudohyphae intertwined with each other, was formed during the temporal development of biofilm, as shown in the hSRS stack image (Fig. 2E). The decomposed SRS images show significant EE accumulation in the fungal biofilm (Fig. 2E). The SRS spectra of lipids and the statistical analysis confirmed that EE accumulated at a high level, which was comparable to the yeast form of *C. albicans* TWO7241 in stationary phase (Fig. 2F and 2G). The acyl C=C level remained markedly high in cells both in stationary phase and biofilm form, which was at higher level compared to cells in the log phase (Fig. 2H). Altogether, these data demonstrate that EE accumulation is a signature of *Candida* biofilm.

### EE in azole-resistant *C. albicans* arises from *de novo* lipogenesis and is largely in the form of ergosteryl oleate

To identify the source of increased EE accumulation in azole-resistant *C. albicans* cells, we examined the contribution of *de novo* lipogenesis and exogenous fatty acid uptake, respectively. Cytosolic acetyl coenzyme A (acetyl-CoA) is the central metabolic intermediate that is essential for lipid biosynthetic reactions through different carbon metabolism pathways, such as glycolysis, β-oxidation, and the glyoxylate cycle (49). Among these metabolic pathways, glucose is universally utilized as the preferred carbon source by most organisms (50, 51). To evaluate the contribution of *de novo* lipogenesis to the increased EE accumulation in azole-resistant *C. albicans* strains, we examined the effects of glycolysis on carbon utilization and lipid storage. Azole-resistant *C. albicans* TWO7241, TWO7243, and azole-susceptible W. Type were cultured in glucose-supplemented medium or glucose-deficient medium until cells reached the stationary phase. The fingerprinting hSRS images of cells grown in glucose-supplemented or glucose-deficient media were acquired, and the hSRS spectra from the LDs were quantified (Fig. 3, A and B, *SI Appendix*, Fig. S7, A and C). We found a significant decrease in the total level of LDs, especially in the accumulation of EE, from cells cultured in glucose-deficient medium compared to glucose-supplemented medium in the azole-resistant TWO7241, TWO7243 strains and the azole-susceptible W. Type strain (Fig. 3C, *SI Appendix*, Fig. S7, B and D). Additionally, the acyl C=C lipid was significantly decreased after glucose depletion (Fig. 3D). This result indicates that glycolysis was a major contributor of the accumulated lipids.

**Fig. 3.**
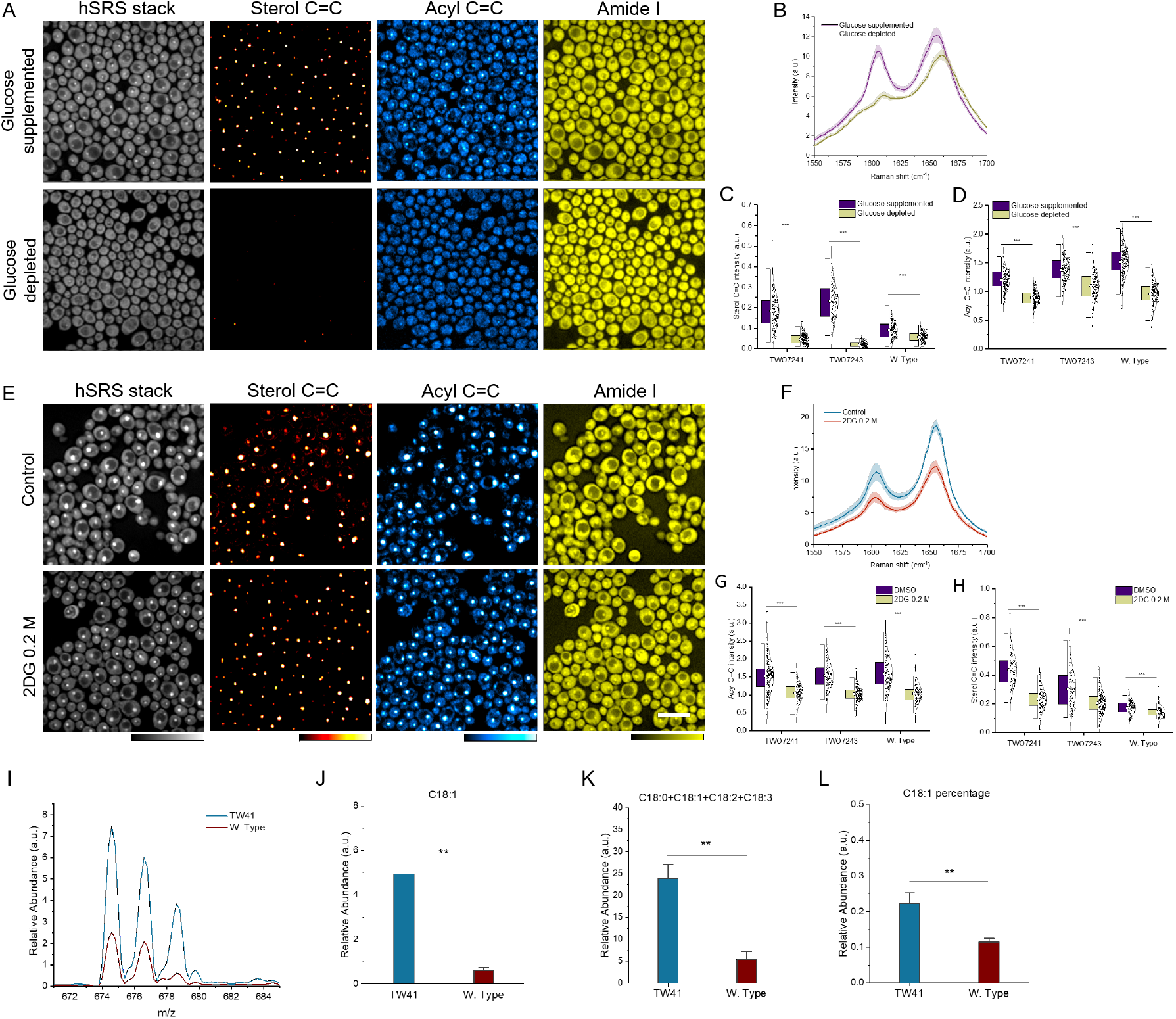
EE accumulation in azole-resistant *C. albicans* cells is related to glucose *de novo* lipogenesis. (A) Spectra unmixing hyperspectral SRS imaging of *C. albicans* cells under glucose depletion treatment. (B) hSRS spectra of lipid accumulation in (A). (C) EE and (D) acyl C=C quantification analyses of hSRS unmixed concentration maps in (A). (E) Spectra unmixing hyperspectral SRS imaging of *C. albicans* cells in the presence of the glycolysis inhibitor (2DG). (F) hSRS spectra of lipid accumulation in (E). (G) EE and (H) acyl C=C quantification analyses of hSRS unmixed concentration maps in (E). (I) Mass spectra of lipids extracted from azole-susceptible *C. albicans* W. Type and azole-resistant *C. albicans* TWO7241 cells. (J) Ergosteryl oleate (EE C18:1) level analysis of mass spectra. (K) Overall lipid level analysis of mass spectra. (L) Quantitative ergosteryl oleate (EE C18:1) to overall lipids (EE C18:0+ C18:1+ C18:2+ C18:3) intensity ratio of *C. albicans* W. Type and *C. albicans* TWO7241 cells. Bar scale represents 10 μm. Significance was evaluated using an unpaired *t*-test (**, p < 0.01; ***, p < 0.001).

Next, we used a glycolysis inhibitor, 2-deoxy-D-glucose (2DG), to further confirm that *de novo* biosynthesis is a major route to the elevated EE storage in azole-resistant *C. albicans*. 2DG, an analog of glucose, cannot undergo further glycolysis since the 2-hydroxyl group in the glucose molecule is replaced by a hydrogen. To assess the effects of 2DG on lipid storage, we first studied its toxicity to fungal cells. The cell viability result under concentration-dependent treatment of 2DG confirmed that the concentration of 0.2 M did not reduce *C. albicans* growth *in vitro* (*SI Appendix*, Fig. S8). The fingerprint hSRS images of cells cultured in YPD medium supplemented with 2DG and cells cultured in normal YPD medium were acquired. As indicated in Fig. 3E and 3F, we observed that EE accumulation was markedly attenuated upon glycolysis inhibition by 2DG in the azole-resistant *C. albicans* TWO7241 and TWO7243 strains. In contrast, upon exposure to 2DG, a less drastic reduction in EE level was observed in azole-sensitive cells compared to fluconazole-resistant cells (Fig. 3G, *SI Appendix*, Fig. S9, A and C). The acyl C=C intensity reduction was not significantly affected in fluconazole-susceptible and fluconazole-resistant cells (Fig. 3H, *SI Appendix*, Fig. S9, B and D). These data together indicate that EE accumulation in azole-resistant *C. albicans* cells is largely due to glucose uptake and *de novo* synthesis. The inhibition of glycolysis effectively reduced the level of EE in the fluconazole-resistant strain.

In order to identify the fatty acid types in the accumulated EE, we performed electrospray ionization mass spectrometry analysis of the extracted lipids from *C. albicans*. Our result revealed that ergosteryl oleate (EE C18:1) accumulated in intracellular lipids was identified to be the dominant species (Fig. 3I). The m/z 679, m/z 677, and m/z 675 peaks correspond to ergosteryl oleate (EE C18:1), ergosteryl linoleate (EE C18:2), and ergosteryl linolenate (EE C18:3), respectively. The quantitative analysis further showed that the level of EE (C18:1) was significantly higher in the TWO7241 strain compared to the wild-type strain (Fig. 3J). Moreover, the total amount of lipids was significantly higher in azole-resistant cells compared to azole-sensitive cells (Fig. 3K). Quantitative analysis showed that the percentage of ergosteryl oleate (EE C18:1) in overall lipids (EE C18:0+ C18:1+ C18:2+ C18:3) is in significant higher level in TWO7241 cells than that in W. Type cells (Fig. 3L).

### Inhibition of EE accumulation by oleic acid effectively impairs azole resistance in stationary-phase *C. albicans* both *in vitro* and *in vivo*

It has been known that sterols are known to be esterified by acyl-CoA-cholesterol acyltransferase (ACAT), which forms steryl esters in an intracellular acyl-CoA-dependent reaction. The two ACAT-related enzymes, Are1p and Are2p, catalyze sterol esterification in yeast (52). The mass data of lipid profiling led to our hypothesis that oleic acid (OA) can be employed as a competitive inhibitor of acyl-CoA to interfere with the active site of the enzyme. This prevents the substrate, acyl-CoA, from binding to the enzyme. To test our hypothesis, we measured whether cell viability or cell growth is affected by oleate treatment. To trace cellular response of OA treatment, we cultured cells in medium supplemented with OA at different concentrations for 13 h and detected the fingerprinting hyperspectral SRS imaging signal as a measurement of exogenous fatty acid uptake. The cell morphology of the azole-resistant strain *C. albicans* TWO7241 was significantly affected by a high concentration of OA treatment (100, and 500 μM), which is indicated by the distorted cell shapes in the transmission images (*SI Appendix*, Fig. S10). In comparison, no morphological changes were observed with *C. albicans* W. Type cells under OA treatment at 10, and 100 μM; morphological changes were not observed until a high concentration of 500 μM OA was used. This result suggests that the cell morphology of azole-resistant *C. albicans* is more vulnerable damage under OA treatment compared to that of azole-sensitive *C. albicans* cells.

To conduct a comprehensive study of the cellular changes in chemical information, we conducted fingerprinting hSRS to inspect metabolic changes in the presence of OA treatment. The hSRS unmixing concentration maps clarified that the intensity of the lipids, including both sterol C=C and acyl C=C, remained at a high level compared to the control and the low dose of OA treatment (10 μM). However, the lipid maps showed almost completely diminished sterol C=C intensity in the presence of a high concentration of OA treatment (100 and 500 μM). The protein signals were remarkably decreased as well, with metabolic heterogeneity observed in the decomposed maps (Fig. 4A). The intensity profile of the lipids showed active EE synthesis in the control and the low dose OA-treated (10 μM) cells, but EE synthesis was dramatically reduced in the presence of higher concentrations of OA (100 and 500 μM) (Fig. 4B). This indicated that the metabolic inhibition in azole-resistant *C. albicans* was visualized by tracing biomass metabolic synthesis under OA treatment. A comparison of the unmixed SRS image intensity further revealed that the synthesis of lipids was highly active in azole-sensitive *C. albicans* cells exposed to 10 μM of OA treatment, but the synthesis of lipids was much reduced in the presence of a higher concentration of OA (100 and 500 μM). However, the unmixing results exhibited a largely diminished signal in the EE image but no significant change in the acyl C=C image until the OA dosage was increased up to 500 μM (*SI Appendix*, Fig. S10). Comparing the unmixed fingerprinting channels between azole-resistant and azole-sensitive *C. albicans* cells, we postulated that the cell viability of azole-resistant *C. albicans* is much more vulnerable to OA treatment, and that EE production is impaired in the presence of OA. To confirm this finding, we investigated cell viability using an optical density measurement. The cell growth of azole-resistant *C. albicans* was not affected in the presence of OA treatment at a low concentration (10 μM). However, OA at 100 and 500 μM effectively inhibited the growth of azole-resistant *C. albicans* cells (Fig. 4D). To ensure that this inhibition was not an acidic effect, we tested the ester form of OA, ethyl oleate (EO), and observed the same concentration-dependent growth inhibition results (Fig. 4E). In contrast, the sensitive species were more robust to OA treatment (*SI Appendix*, Fig. S10). In summary, our observation supports that EE inhibition by OA reduces the viability of azole-resistant fungi.

**Fig. 4.**
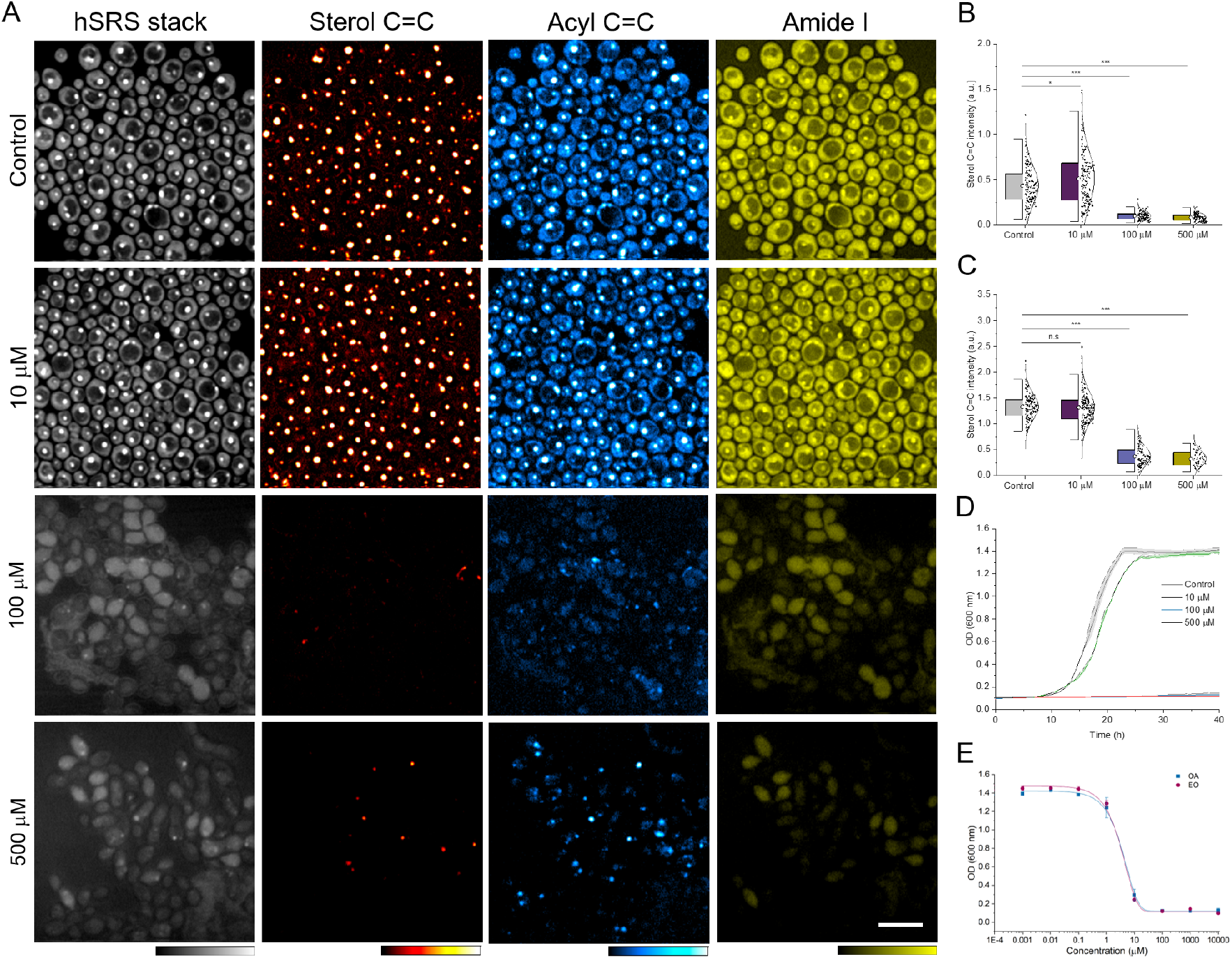
Oleate attenuates EE accumulation in azole-resistant *C. albicans*. (A) Fingerprinting hyperspectral SRS images of stationary-phase *C. albicans* cells under concentration-dependent oleate treatment. Quantification of (B) EE and (C) acyl C=C levels in hSRS unmixed concentration maps to show OA inhibition on EE accumulation. (D) Growth inhibition of *C. albicans* TWO7241 under concentration-dependent oleate treatment. (E) Comparison of growth inhibition of *C. albicans* TWO7241 under concentration-dependent OA and EO treatment. Bar scale represents 10 μm. Significance was measured using an unpaired *t*-test (*, p < 0.05; ***, p < 0.001; n.s., not significant).

Because ergosterol esterification is known to play a vital role in maintaining intracellular ergosterol homeostasis, we evaluated how cell susceptibility to azole antifungals could be affected by oleate-mediated abrogation of EE. Additionally, we evaluated whether the combination of OA and azoles would exhibit a synergistic relationship and reduce azole tolerance in fungi. To determine if a synergistic relationship exists, we used the checkerboard assay to monitor the optical density of azole-resistant *C. albicans* TWO7241, TWO7243, NR-29446, ATCC 64124 in the presence of oleate and fluconazole treatment. A synergistic relationship was identified between oleate and fluconazole treatment against azole-resistant *C. albicans* (Fig. 5A). Notably, the lowest azole concentration that inhibited *C. albicans* TWO7241 growth within 24 h steadily decreased when the dose of OA was increased. An OA dose of 128 μg/ml resulted in a 16-fold reduction in the minimum inhibitory concentration (MIC) of fluconazole, where a two-fold change or larger was classified as synergy based on the fractional inhibitory concentration index (FICI). A synergistic relationship was also observed between OA and fluconazole against other azole-resistant strains: for *C. albicans* TWO7243, a 16-fold reduction in the MIC of fluconazole was observed in the presence of 64 μg/ml OA; for *C. albicans* NR-29446, a 64-fold reduction in the MIC of fluconazole was observed in the presence of 128 μg/ml OA; for *C. albicans* ATCC 64124, a 16-fold reduction in the MIC of fluconazole was observed in the presence of 128 μg/ml of OA treatment, respectively. The calculation of FICI based on the MICs of fluconazole or OA and the fractional inhibitory concentration (FIC), confirmed the synergistic effect between oleate and fluconazole (Fig. 5B). Additionally, the combination of fluconazole (at 8 μg/ml) with OA, at concentrations of 10 μM and higher, reduced the growth of *C. albicans* as observed over a 48-hr period (Fig. 5C). The results confirmed that using OA, an EE biosynthesis inhibitor, significantly impaired the cell viability and resistance to fluconazole in azole-resistant *C. albicans* strains. These growth inhibition results further validate our hypothesis that OA with fluconazole exhibits a strong synergistic effect in suppressing the growth of azole-resistant *C. albicans* cells compared to either agent alone. Notably, palmitic acid (PA) and arachidonic acid (AA) did not exhibit a synergistic relationship with fluconazole against *C. albicans* TWO7241 (*SI Appendix*, Fig. S12). OA treatment inhibits ergosterol esterification biosynthesis which is vital for ergosterol homeostasis. The azole antifungals inhibit the ergosterol biosynthesis pathway. Thus, OA acted synergistically with fluconazole against azole-resistant *C. albicans*.

**Fig. 5.**
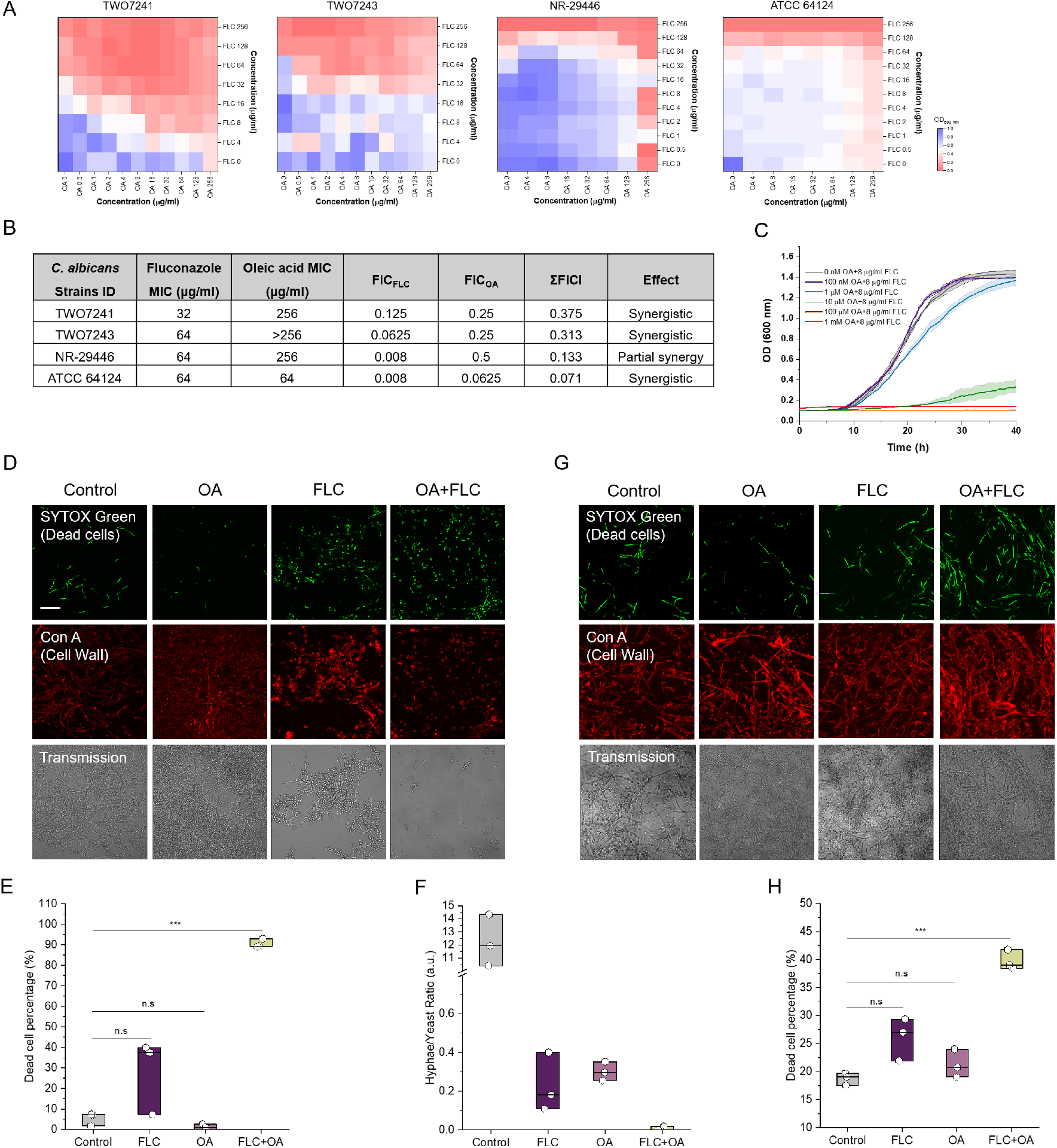
Oleate and fluconazole exhibit a synergistic relationship against azole-resistant *C. albicans* in stationary phase and biofilm development. (A) A synergistic relationship between oleate and fluconazole (FLC) was determined by azole-resistant *C. albicans* strains TWO7241, TWO7243, NR-29446, ATCC 64124, and ATCC MYA573. (B) Fractional inhibitory concentration (FIC) of fluconazole with oleic acid (OA) treatment in azole-resistant *C. albicans* strains TWO7241, TWO7243, NR-29446, ATCC 64124, ATCC MYA573. (C) Growth inhibition of *C. albicans* TWO7241 in the presence of oleate and fluconazole combination treatment. (D) Live and dead assay of OA/FLC synergistic treatment on *C. albicans* to elucidate the inhibition of biofilm formation. The fluorescent green and red signals indicate SYTOX (cell nucleus) and Con A (cell wall), respectively. (E) Synergistic effect of impairing azole tolerance and cell viability through combination therapy with OA/FLC. (F) A histogram that shows a lower hyphae to yeast form ratio under the OA/FLC combination treatment, which indicates the combination induces an inhibitory effect on fungal biofilm development. (G) Live and dead assay of OA/FLC synergistic treatment on the viability of cells in the *C. albicans* biofilm. (H) The quantitative dead cell ratios indicate a substantial suppression of cells in biofilm viability was achieved by OA/FLC combination therapy. Bar scale represents 10 μm. Significance was measured using an unpaired *t*-test (***, p < 0.001; n.s., not significant).

After confirming the efficacy of OA and fluconazole combination treatment in azole-resistant *C. albicans* cells, and the hSRS imaging predicted that inhibition in EE accumulation may impair the biofilm-forming ability of *C. albicans*, we further explored the synergistic effect of OA with fluconazole on the growth of *C. albicans* biofilm. Biofilm development from yeast cells and the biofilm cell viability were examined under OA and azole treatment. The concentration of OA at 128 μg/ml and fluconazole at 16 μg/ml were chosen for combination therapy against *C. albicans* TWO7241 biofilm. We performed confocal fluorescence imaging to identify the dead fungal cells with SYTOX green nucleic acid stain and the overall fungal cells using the cell wall stain Concanavalin A (Con A) as an indicator. Stationary phase *C. albicans* TWO7241 cells were seeded to grow a biofilm over 24 hours. The OA or azole treatment were then applied before the biofilm was developed from the yeast form of *C. albicans*. As shown in Fig. 5D, the transmission images clearly show that in the control group, cells developed large number of filamentous hyphae with extracellular matrix after the 24-hour incubation period indicating a biofilm had formed. In the OA treatment group, the cells developed large numbers of filamentous hyphae. However, in the FLC or the OA/FLC treatment groups, there was a reduced number of filamentous hyphae and more yeast cells remained. From the green channels showing the dead fungal cells with SYTOX green, we observed that the dead cells ratio was dramatically higher in the OA/FLC group. Quantification of the dead cell (green channel) and the total cell amount (red channel) further confirmed that the ratio of dead cells was significantly higher in response to OA/FLC treatment compared to the other 3 groups (Fig. 5E). This validated the synergistic effect of OA/FLC to impair azole tolerance and cell viability. From the total cell amount indicated in the red channel, we estimated the cell number of yeast form and hyphae form. As expected, the histogram showed the fungal cells largely remained in the yeast form in the presence of OA/FLC, which indicated there was an inhibitory effect in fungal biofilm development (Fig. 5F).

We further evaluated if the OA/FLC combination could eradicate a fungal biofilm. The biofilm of *C. albicans* TWO7241 yeast cells was first grown from for 12 h, and then the OA or FLC treatment was incubated with the biofilm for another 12 h. No signs of morphological changes were detected between the treatment groups (Fig. 5G). The live/dead fluorescence imaging suggested that OA or FLC treatment alone did not affect cell viability over the treatment period. Interestingly, the ratio of dead cells was markedly increased in the presence of the OA/FLC combination when compared to OA or FLC alone. This indicates OA/FLC substantially suppressed the formation of *C. albicans* biofilm. The quantitative ratio of dead cells was calculated and plotted in Fig. 5H. These data demonstrate an enhanced effect of OA and FLC when administrated together to enhance the activity of fluconazole in the biofilm of *C. albicans*.

To evaluate the efficacy of combining OA and FLC to overcome azole resistance *in vivo*, we investigated the effect of OA/FLC in a murine skin wound infection model (53). To induce skin lesions in mice (4 groups [n=2 mice/group]), a fungal suspension containing approximately 10^8^ CFU/ml of azole-resistant *C. albicans* TWO7241 was inoculated on the wounds and uniformly applied gently onto the mice skin (Fig. 6A). Three hours after the wounds were infected, the first topical treatments were administered to each group (FLC at 32 μg/ml or OA at 256 μg/ml). The second treatment was administered 21 hours after the wounds were infected. The wounds of all the treated groups and the control group are shown in Fig. 6B. Then, mice were humanely euthanized, and the wound tissues were aseptically collected in order to quantify the *Candida* filamentation in wounds. Periodic acid–Schiff (PAS) staining was further employed to examine the physiological condition of the wounds. The untreated, OA-treated, and FLC-treated groups all showed the formation of *C. albicans* filaments below the wound, in which dead tissues, yeast or hyphae form fungi, macrophages, and neutrophils dwell (Fig. 6C). This suggests that the immune system of mice fought against fungi residing inside the wound tissue. Treatment of OA alone did not significantly influence *C. albicans* hyphae development relative to the untreated control (p > 0.05) (Fig. 6C, 6D). In contrast, OA/FLC effectively inhibited the formation of *C. albicans* hyphae in mice skin tissues, with yeast from *C. albicans* aggregated on the mice skin surface. These results qualitatively and quantitatively demonstrate the improvement of OA/FLC in their ability to impair *Candida* filamentation *in vivo* (Fig. 6C, 6D). The synergistic relationship between OA and FLC, as demonstrated here, implies a novel approach to effectively inhibit the growth of *C. albicans* hyphae, which impairs biofilm formation.

**Fig. 6.**
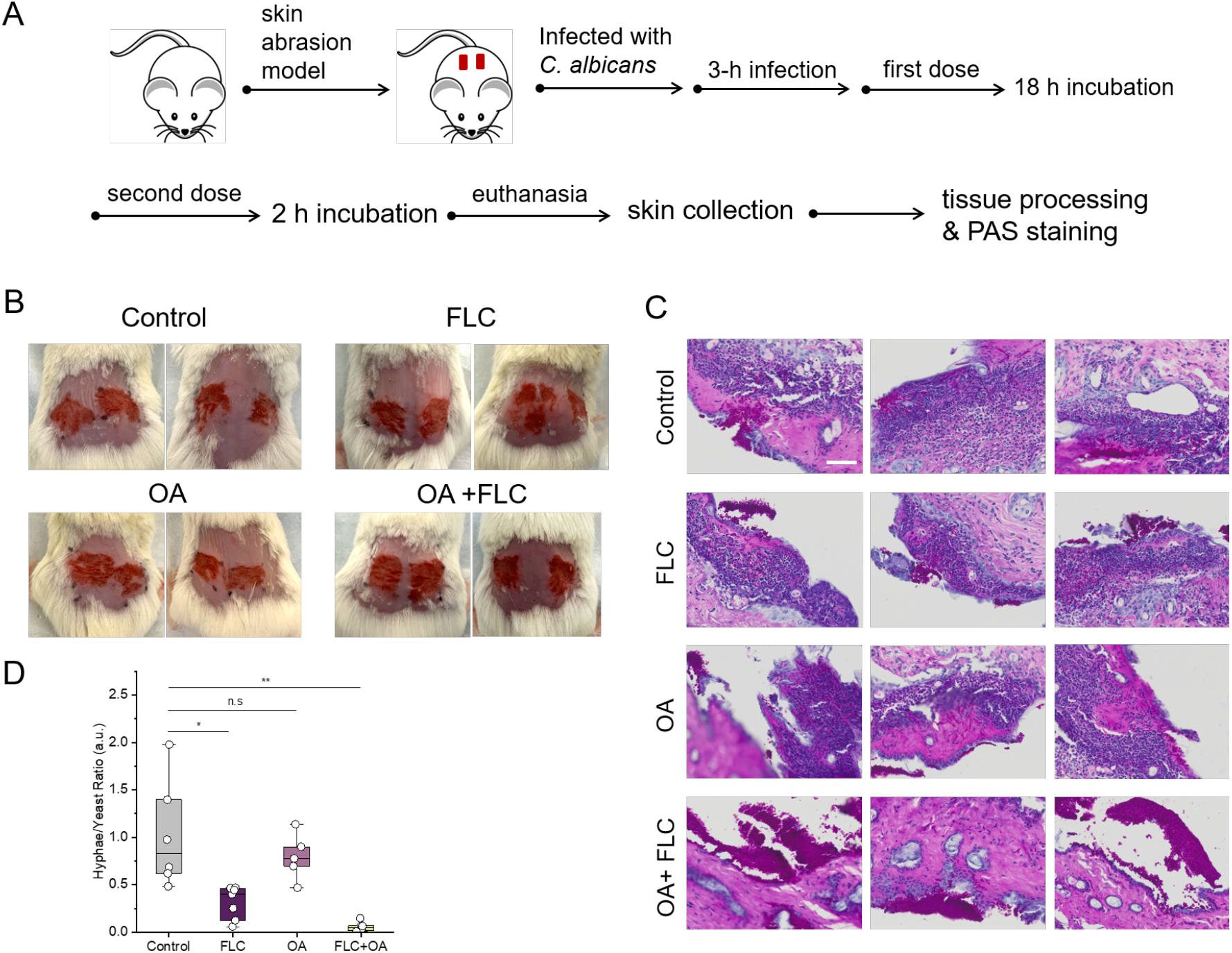
Inhibition of EE accumulation by oleic acid effectively impairs azole resistance of *C. albicans in vivo*. (A) Schematic illustration of development and subsequent treatment for *C. albicans*-induced mice skin abrasions. (B) Pictures of murine skin wounds of 4 different groups taken before treatment; (C) Histology scanning of PAS staining of *C. albicans*-infected murine skin tissue in the presence of different treatments. Bar scale represents 50 μm. (D) Ratio of hyphae to yeast cells after *C. albicans* was exposed to different treatments in (A). Significance was measured using an unpaired *t-*test (*, p < 0.05; **, p < 0.01; n.s., not significant).

## Discussion

Multidrug-resistant *Candida* species are rapidly emerging and spreading globally. The mortality rates of invasive *C. albicans* infection remain high despite the availability of existing antifungal therapies. Strategies that can combat the emergence and spread of antifungal resistance are crucial for guiding therapeutic treatment. However, an understanding of the underlying mechanism of fungal cell metabolism reprogramming in response to azole treatment is incomplete. *C. albicans* is capable of forming highly drug-resistant biofilms, an organized three-dimensional structure that is comprised of a dense network of cells in an extracellular matrix of carbohydrates, glycoproteins, lipids, and nucleic acids (54-56). These biofilms restrict access to echinocandin drugs, and they are intrinsically resistant to azoles (10, 57). As the biofilms of *C. albicans* are recalcitrant to antifungal treatment, biofilms pose a significant medical challenge for the treatment of candidiasis. (58) The development and formation of biofilms is a multi-step process that involves various adaptive mechanisms, such as lipid composition alteration (47). Cells in *C. albicans* biofilms undergo phase-dependent changes in the levels and composition of lipids (59, 60).

Here, by hyperspectral SRS imaging that enables visualization and quantitative analysis of lipid metabolism integrated with LASSO analysis to quantify the intracellular chemical contents, we report an aberrant accumulation of EE in azole-resistant *C. albicans* as compared with non-resistant species at a single cell level. Such accumulation is found to arise from *de novo* glucose lipogenesis. According to lipid profiling analysis by mass spectrometry, ergosterol oleate storage significantly increases in azole-resistant *C. albicans*. Consequently, blocking EE accumulation by using azoles in combination with oleate synergistically suppressed *C. albicans* cell viability *in vitro* and the growth of biofilms on the wounds of mice *in vivo*.

Visualizing metabolism in single living cells has been challenging due to technical difficulties. Here, by fingerprinting hSRS imaging, we demonstrated visualization and quantitative analysis of lipid metabolism at the single cell level in a temporal and spatially resolved manner. This method is complementary to current techniques, like mass spectrometry, nuclear magnetic resonance spectroscopy, fluorescence imaging, or single-color SRS spectroscopy. Instead of ensemble measurement, high spatial resolution is vital for exploring intracellular dynamic and complex metabolic networks. Visualizing the mechanisms underlying fungal resistance to azole antifungals and revealing the metabolic heterogeneity or the diversity in metabolism at a single cell level should facilitate a better understanding of why some fungal species are intrinsically resistant to azoles. Our method opens an avenue to address this question by imaging the metabolic response in a wide variety of fungal cells or a biofilm *in situ*. Another important question that can be pursued by our technology is whether a therapeutic strategy can be developed through a quantitative, comparative study of intracellular metabolites between sensitive fungal cells, resistant cells, and biofilm cells.

In this work, we showed that compared to azole-sensitive *C. albicans* cells, resistant cells exhibit significantly higher level of EE accumulation derived mainly from *de novo* glucose lipogenesis. Our observation is consistent with previous reports of higher EE accumulation levels in some azole-resistant cells (61, 62). A recent study reported significant enrichment of genes associated with ergosterol and sphingolipid biosynthesis in fluconazole-treated cells which has the highest correlation with fluconazole resistance (63). Oleate inhibited steryl ester synthesis and caused liposensitivity in yeast (64). However, direct evidence to elucidate azole resistance and steryl esterification is needed. The difference of EE biosynthetic preference for *C. albicans* may be related to its special metabolic demands, leading to our observation of a distinct EE biosynthetic metabolic pathway in azole-resistant *C. albicans*. We also noticed that different clinical isolates may have distinct metabolic profiles. Azole-resistant *C. albicans* strains ATCC MYA573, ATCC 64124, and NR-29448 showed relatively lower cellular levels of EE. Further investigation is needed to fully understand the metabolic networks on how high cellular levels of EE contribute to azole resistance. Our imaging method could be a powerful tool to reveal the metabolic differences between different cell models in clinically resistant isolates and other fungal pathogens. Developing more applications for our approach relies on improving imaging sensitivity further. Due to the current limited detection sensitivity at a millimolar level, we could not detect ergosterol or sphingolipid on the cell membrane in the fingerprint region. Higher sensitivity would allow mapping of the complex organization with distinct lipid compositions on cell membranes. Future elucidation is needed of the molecular mechanisms by which the EE biosynthetic pathway will determine whether ergosterol esterification is a compelling therapeutic target across multiple *Candida* types. Regulating ergosterol metabolism in *Candida* cells from multiple isolates will further improve the current understanding of how metabolic transformation is linked to antifungal resistance.

## Materials and Methods

### *C. albicans* clinical isolates and antifungal susceptibility testing

A wide variety of *C. albicans* strains with different levels of susceptibility to fluconazole were utilized and described in (*SI Appendix*, Table S1). The list includes 10 fluconazole-sensitive strains (MIC ≤ 2 μg/ml), 6 fluconazole-resistant strains (MIC ≥ 8 μg/ml), and 4 fluconazole-susceptible dose-dependent strains (MIC 2-4 μg/ml).

### Chemicals and reagents

Yeast extract peptone dextrose (YPD), yeast nitrogen base (YNB), yeast extract, 2-deoxy-D-glucose (2DG), thiazolyl blue tetrazolium blue (MTT), and oleic acid were purchased from Sigma-Aldrich (St. Louis, MO). Yeast extract peptone (YP) medium was prepared by adding 20 g of bacteriological peptone (Becton, Dixon, Franklin, NJ) and 10 g of yeast extract in 1 l of distilled water. Phosphate-buffered saline (PBS) and fluconazole were purchased from Thermo Fisher Scientific (Waltham, MA).

### Cell culture conditions

*C. albicans* isolates were initially cultured by inoculating a single colony in sterile YPD broth at 30°C in an orbital shaker (VWR, model 3500I) at a shaking speed of 200 rpm at a tilted angle of 45°. Stationary-phase cells are harvested after 24 h of incubation. Logarithmic-phase cells were prepared by 1:20 dilution of an overnight culture of fungi into fresh pre-warm YPD and cultured for another 5 to 6 h. Cells were then centrifuged. The supernatant was discarded and then resuspended and washed with fresh PBS. Next, fungal cells were fixed in 10% formalin before the formalin was washed away twice using 1× PBS before imaging. The fungal specimen was sandwiched between 2 cover glasses (VWR international).

For the glucose depletion study, cells in the exponential phase were cultured in YPD and YP medium overnight. For the glycolysis inhibition assay, cells in the exponential phase were cultivated in YPD and YPD supplemented with 0.2 M 2DG for 13 h.

In the glycolysis inhibition toxicity test, cells were grown overnight in YPD. Cells were then washed three times in PBS and optical density (OD) adjusted to 0.5 at 600 nm. The cells were then inoculated in YPD supplemented with 0.4 M 2DG. Two hundred microliters of the cells with 0.4 M 2DG in YPD medium were seeded onto a 96-well plate. A two-fold serial dilution to a final concentration of 0.025 M 2DG was performed. The cells were then incubated at 30 °C for 13 h. Cell viability was measured with the PrestoBlue cell viability assay (Thermo Fisher).

### SRS imaging

Hyperspectral SRS imaging was conducted with a spectral focusing method (65, 66). Briefly, the Raman shift was tuned by controlling the temporal delay between 2 chirped femtosecond pulses. A femtosecond laser (Coherent) operating at 80 MHz provided the pump and Stokes laser source. With the pump beam tuned to 891 nm, Stokes beam was tuned to 1040 nm to cover the fingerprint C=C vibrational region. The Stokes beam was modulated at 2.3 MHz by an acousto-optic modulator (1205-C, Isomet). After combination, both the pump and Stokes beams were chirped by 12.7 cm long SF57 glass rods and then sent to a laser-scanning microscope. A 60x water immersion objective (NA = 1.2, UPlanApo/IR, Olympus) was used to focus the light on the sample. An oil condenser (NA = 1.4, U-AAC, Olympus) was used to collect the signal.

To acquire hyperspectral SRS images, a stack of 120 images at different pump-Stokes temporal delay was recorded. The temporal delay was controlled by an automatic stage that moved forward with a step size of 10 μm. To calibrate the Raman shift to the temporal delay, standard chemicals, including DMSO, triglyceride, and ergosterol, with known Raman peaks in C=C region from 1460 to 1750 cm^−1^ were used. The average acquisition time for a 200 × 200 pixels image was about 1 second. Hyperspectral SRS images were analyzed using ImageJ (National Institute of Health).

### Spontaneous Raman spectroscopy

Confocal Raman spectral analysis from individual fungal cells was performed. An excitation laser at 532 nm was used for Raman spectral acquisition. The acquisition time for a typical spectrum from individual fungal cells was 20 seconds, with the beam power maintained around 10 mW at the sample. For each specimen, at least 10 spectra from different individual cells were obtained. The background was removed manually, and the peak height was measured.

### Electrospray ionization mass spectrometry (ESI-MS) measurement of lipid extraction

Lipid extraction from cell pellets was performed according to Folch et al (67). Stationary-phase *C. albicans* cells were collected and counted. The extraction of lipids was conducted by mixing large pellets to 300 μl of water/vial and homogenized well with vortexing. CK5 precellys tubes were used to disrupt fungal walls for B&D extraction. The dried sample was diluted in 50 ml of 1:1 CHCl3:MeOH, 10 mM NH4Ac. The cell suspension (20 ml) was mixed with 20 mL of internal standard solution (300 ng/μl of C17-cholesteryl ester (CE), molecular weight = 656 g/mol). ESI-MS analysis was conducted according to the protocol described previously (68) using 2 precursor ion scans with CE using 369.3 (m/z) and ergosterols using 379.3 (m/z). The relative level of ergosteryl oleate (18:1) was normalized by cell number for comparison between sensitive and resistant cells.

### Fluorescence imaging of live and dead *C. albicans* biofilm

An overnight culture of *C. albicans* TWO7241 was grown in a 30 °C incubator with shaking (at 250 rpm). The overnight culture of *C. albicans* was diluted (1:100) in RPMI-1640 and transferred to glass bottom dishes (35 mm, In Vitro Scientific). The plates were incubated at 37 °C with supplementary 5% CO_2_ for 12 to 24 h to form mature biofilm. Thereafter, the media was removed and the surface of the dish was washed gently with PBS to remove planktonic fungal cells. Plates were subsequently treated with fluconazole, OA, or a combination of fluconazole and OA. After treatment, biofilms were immediately stained with fluorescent dyes. To confirm the existence of biofilm on the glass bottom surface, the biofilm matrix was stained by Con A (Invitrogen). To quantify dead cells, biofilms are incubated with Sytox Green, a cell-impermeable fluorescent DNA dye to quantify the survival percent of *C. albicans* in the biofilm after treatment. The biofilms were washed with PBS twice and then imaged using a fluorescence microscope (OLYMPUS FV3000, objective: 40X, oil immersion, NA = 1.3). Two different excitation channels (live: excitation at 488 nm; dead: excitation at 561 nm) were utilized in order to map the ratio of live versus dead cells within the biofilm. The acquired images were analyzed by ImageJ (National Institute of Health).

### Checkerboard broth dilution assays

To evaluate the combinatorial behavior between azole and oleate, we performed checkerboard broth dilution assays to calculate the fractional inhibition centration index. Logarithmic-phase *C. albicans* inoculum were transferred to a 96-well plate containing two-fold dilution of OA starting at 256 μg/ml, followed by two-fold dilution of fluconazole starting at 256 μg/ml in a perpendicular direction. Then the plate was cultured at 30 °C for 24 h. The optical density at 600 nm (OD 600 nm) was recorded to represent the fungal cell number. A heat map correlated with OD 600 nm was generated to calculate the FICI.

### *In vivo* assessment of synergy between fluconazole and OA

This study was approved by the Institutional Animal Care and Use Committee at Boston University and was performed in the Animal Science Center in the Charles River Campus. The abrasion fungal infection model was used. Eight BALB/c mice (Jackson Laboratories, 000651) aged 6–8 weeks and weighing about 20 g were placed under anesthesia and shaved on the dorsal side before producing the abrasion wounds. The tissue was carefully abraded within a defined 1.0 cm × 1.0 cm area using a #15 sterile scalpel blade. The scraped area did not produce any blood out of the skin barrier. After producing the abrasion, 50 μl of fungal suspension containing approximately 10^8^ CFU/ml of stationary phase *C. albicans* TWO7241 in PBS was inoculated onto the abrasion wound and uniformly applied gently using the side of a pipette tip and air dried. Three hours after inoculating the fungal suspension, OA or FLC treatment was topically applied on the infected wounds. Treatments were applied to mice twice, the first treatment being applied 3 h following infection, and the second treatment was applied 21 h after the infection. Two h after the second treatment, mice were euthanized. The wound tissue was harvested and fixation by 10% neutral buffered formalin for periodic acid-Schiff staining and histology studies. The tissue histology images were acquired using a slide scanning microscope (OLYMPUS VS120).

### Spectral unmixing and single cell analysis

To decompose the hSRS stack into chemical maps, the spectral unmixing for was performed by applying a pixel-wise least absolute shrinkage and selection operator (LASSO) regression algorithm. Quantification of droplet size and intensity in individual fungal cells was done by applying automatic local thresholding and analyzing particles to the hSRS image stack. Data analysis and visualization were done using CellProfiler (*SI Appendix*, Fig. S4). The Student’s unpaired *t-*test was used to determine whether there was any statistically significant difference between treatment groups (*, p < 0.05; **, p < 0.01; ***, p < 0.001; n.s., not significant.).

## Acknowledgments

This work is supported by R01 AI141439 and R35 GM136223. We would like to thank Dr. Theodor White (University of Missouri-Kansas City), Dr. David Rogers (University of Tennessee Health Science Center), and BEI resources for kindly providing *C. albicans* isolates used in this study. Research reported in this publication was supported by the Boston University Micro and Nano Imaging Facility and the Office of the Director, National Institutes of Health under award Number S10OD024993. The content is solely the responsibility of the authors and does not necessarily represent the official views of the National Institute of Health. We acknowledge Christina R. Ferreira and Bruce R. Cooper from Purdue Metabolomics Facility for their help on mass spectrometry measurements. We would like to thank Fukai Chen for the help on histology slides scanning assay.

## Author Contributions

M. Z. and J.-X. C conceived the idea. M. N. S. provided the clinical fungal isolates and constructive discussions. M. Z., P.-T.D., J.-X. C., and M. N. S. designed the experiments. M. Z. and P.-T. D. designed, performed and analyzed initial SRS and fluorescence imaging experiments. M. Z. designed, performed, and analyzed *in vitro* mechanism studies and synergistic therapy studies. M. Z. designed and performed the biofilm growth assays, fluorescence assays, and imaging experiments. Y. Z. conducted the *in vivo* mice abrasion experiments. H. E. and E. S. performed the MIC assay, checkerboard assay and interpretation, helped with biofilm experiments. Y. Z. helped with the histology slides scanning assay. H. L. developed the hyperspectral SRS unmixing method. Z. W. participated in part of the cell imaging data analysis. S. J. helped with the *in vivo* studies. C. Z. helped with the SRS imaging measurements and data analysis. H. E., E. S., and Z. C. provided constructive suggestions over the project and manuscript. J.-X. C. supervised the overall project. M. Z. and J.-X. C. co-wrote the manuscript. M. N. S. revised the manuscript. All authors read and commented on the manuscript.

## References

1. W. Denning David, J. Bromley Michael, How to bolster the antifungal pipeline. Science 347, 1414–1416 (2015).

2. D. Brown Gordon et al., Hidden Killers: Human Fungal Infections. Sci. Transl. Med. 4, 165rv113–165rv113 (2012).

3. M. Kojic Erna, O. Darouiche Rabih, Candida Infections of Medical Devices. Clin. Microbiol. Rev. 17, 255–267 (2004).

4. M. A. Pfaller, D. J. Diekema, Epidemiology of Invasive Mycoses in North America. Crit. Rev. Microbiol. 36, 1–53 (2010).

5. M.-F. Cheng et al., Risk factors for fatal candidemia caused by Candida albicans and non-albicans Candida species. BMC Infect. Dis. 5, 22 (2005).

6. M. Morrell, J. Fraser Victoria, H. Kollef Marin, Delaying the Empiric Treatment of Candida Bloodstream Infection until Positive Blood Culture Results Are Obtained: a Potential Risk Factor for Hospital Mortality. Antimicrob. Agents Chemother. 49, 3640–3645 (2005).

7. K. C. Howard, E. K. Dennis, D. S. Watt, S. Garneau-Tsodikova, A comprehensive overview of the medicinal chemistry of antifungal drugs: perspectives and promise. Chem. Soc. Rev. 49, 2426–2480 (2020).

8. J. E. Nett, D. R. Andes, Antifungal Agents: Spectrum of Activity, Pharmacology, and Clinical Indications. Infectious Disease Clinics of North America 30, 51–83 (2016).

9. P. G. Pappas et al., Clinical Practice Guideline for the Management of Candidiasis: 2016 Update by the Infectious Diseases Society of America. Clin. Infect. Dis. 62, e1–e50 (2015).

10. D. S. Perlin, E. Shor, Y. Zhao, Update on Antifungal Drug Resistance. Curr. Clin. Microbiol. Rep. 2, 84–95 (2015).

11. S. G. Whaley et al., Azole Antifungal Resistance in Candida albicans and Emerging Non-albicans Candida Species. Front. Microbiol. 7 (2017).

12. P. Marichal et al., Contribution of mutations in the cytochrome P450 14α-demethylase (Erg11p, Cyp51p) to azole resistance in Candida albicans. Microbiology 145, 2701–2713 (1999).

13. T. Coste Alix, M. Karababa, F. Ischer, J. Bille, D. Sanglard, TAC1, Transcriptional Activator of CDR Genes, Is a New Transcription Factor Involved in the Regulation of Candida albicans ABC Transporters CDR1 and CDR2. Eukaryot. Cell 3, 1639–1652 (2004).

14. T. Liu Teresa et al., Genome-Wide Expression and Location Analyses of the Candida albicans Tac1p Regulon. Eukaryot. Cell 6, 2122–2138 (2007).

15. A. Coste et al., Genotypic Evolution of Azole Resistance Mechanisms in Sequential Candida albicans Isolates. Eukaryot. Cell 6, 1889–1904 (2007).

16. S. L. Kelly, D. C. Lamb, D. E. Kelly, Sterol 22-desaturase, cytochrome P45061, possesses activity in xenobiotic metabolism. FEBS Lett. 412, 233–235 (1997).

17. F. S. Nolte et al., Isolation and characterization of fluconazole- and amphotericin B-resistant Candida albicans from blood of two patients with leukemia. Antimicrob. Agents Chemother. 41, 196–199 (1997).

18. Y. Miyazaki et al., Cloning, sequencing, expression and allelic sequence diversity of ERG3 (C-5 sterol desaturase gene) in Candida albicans. Gene 236, 43–51 (1999).

19. S. Chau Andrew et al., Inactivation of Sterol Δ5,6-Desaturase Attenuates Virulence in Candida albicans. Antimicrob. Agents Chemother. 49, 3646–3651 (2005).

20. M. Martel Claire et al., Identification and Characterization of Four Azole-Resistant erg3 Mutants of Candida albicans. Antimicrob. Agents Chemother. 54, 4527–4533 (2010).

21. F. Morio, F. Pagniez, C. Lacroix, M. Miegeville, P. Le Pape, Amino acid substitutions in the Candida albicans sterol Δ5,6-desaturase (Erg3p) confer azole resistance: characterization of two novel mutants with impaired virulence. J. Antimicrob. Chemother. 67, 2131–2138 (2012).

22. L. E. Cowen, D. Sanglard, S. J. Howard, P. D. Rogers, D. S. Perlin, Mechanisms of antifungal drug resistance. Cold Spring Harb. Perspect. Med. 5, a019752 (2015).

23. S. Yue et al., Cholesteryl ester accumulation induced by PTEN loss and PI3K/AKT activation underlies human prostate cancer aggressiveness. Cell Metab 19, 393–406 (2014).

24. J. Li et al., Lipid Desaturation Is a Metabolic Marker and Therapeutic Target of Ovarian Cancer Stem Cells. Cell Stem Cell 20, 303-314.e305 (2017).

25. H. J. Lee et al., Multimodal Metabolic Imaging Reveals Pigment Reduction and Lipid Accumulation in Metastatic Melanoma. BME Frontiers 2021, 9860123 (2021).

26. J. Du et al., Raman-guided subcellular pharmaco-metabolomics for metastatic melanoma cells. Nat. Commun. 11, 4830 (2020).

27. F.-K. Lu et al., Label-Free Neurosurgical Pathology with Stimulated Raman Imaging. Cancer Res. 76, 3451–3462 (2016).

28. F.-K. Lu et al., Label-free DNA imaging in vivo with stimulated Raman scattering microscopy. Proc. Natl. Acad. Sci. U.S.A 112, 11624–11629 (2015).

29. W.-W. Chen et al., Spectroscopic coherent Raman imaging of Caenorhabditis elegans reveals lipid particle diversity. Nat. Chem. Biol. 16, 1087–1095 (2020).

30. M. C. Wang, E. J. O’Rourke, G. Ruvkun, Fat Metabolism Links Germline Stem Cells and Longevity in C. elegans. Science 322, 957–960 (2008).

31. L. Shi et al., Optical imaging of metabolic dynamics in animals. Nat. Commun. 9, 2995–2995 (2018).

32. A. J. Chen et al., Fingerprint Stimulated Raman Scattering Imaging Reveals Retinoid Coupling Lipid Metabolism and Survival. ChemPhysChem 19, 2500–2506 (2018).

33. P. Wang et al., Label-free quantitative imaging of cholesterol in intact tissues by hyperspectral stimulated Raman scattering microscopy. Angew. Chem., Int. Ed. Engl. 52, 13042–13046 (2013).

34. P. Wang et al., Imaging Lipid Metabolism in Live Caenorhabditis elegans Using Fingerprint Vibrations. Angew. Chem. Int. Ed. 53, 11787–11792 (2014).

35. H. J. Lee et al., Assessing Cholesterol Storage in Live Cells and C. elegans by Stimulated Raman Scattering Imaging of Phenyl-Diyne Cholesterol. Sci. Rep. 5, 7930 (2015).

36. L. Wei et al., Super-multiplex vibrational imaging. Nature 544, 465–470 (2017).

37. L. Wei et al., Live-cell imaging of alkyne-tagged small biomolecules by stimulated Raman scattering. Nat. Methods 11, 410–412 (2014).

38. F. Hu, L. Shi, W. Min, Biological imaging of chemical bonds by stimulated Raman scattering microscopy. Nat. Methods 16, 830–842 (2019).

39. P.-T. Dong et al., Polarization-sensitive stimulated Raman scattering imaging resolves amphotericin B orientation in Candida membrane. Sci. Adv. 7, eabd5230 (2021).

40. M. Zhang et al., Rapid Determination of Antimicrobial Susceptibility by Stimulated Raman Scattering Imaging of D_2_O Metabolic Incorporation in a Single Bacterium. Adv. Sci. 7, 2001452 (2020).

41. W. Hong et al., Antibiotic Susceptibility Determination within One Cell Cycle at Single-Bacterium Level by Stimulated Raman Metabolic Imaging. Anal. Chem. 90, 3737–3743 (2018).

42. C. W. Karanja et al., Stimulated Raman Imaging Reveals Aberrant Lipogenesis as a Metabolic Marker for Azole-Resistant Candida albicans. Anal. Chem. 89, 9822–9829 (2017).

43. A. Flowers Stephanie et al., Gain-of-Function Mutations in UPC2 Are a Frequent Cause of ERG11 Upregulation in Azole-Resistant Clinical Isolates of Candida albicans. Eukaryot. Cell 11, 1289–1299 (2012).

44. M. Werner-Washburne, E. L. Braun, M. E. Crawford, V. M. Peck, Stationary phase in Saccharomyces cerevisiae. Mol. Microbiol. 19, 1159–1166 (1996).

45. M. Werner-Washburne, E. Braun, G. C. Johnston, R. A. Singer, Stationary phase in the yeast Saccharomyces cerevisiae. Microbiol. Rev. 57, 383–401 (1993).

46. F. R. Taylor, L. W. Parks, Metabolic interconversion of free sterols and steryl esters in Saccharomyces cerevisiae. J. Bacteriol. 136, 531–537 (1978).

47. D. Alim, S. Sircaik, S. L. Panwar, The Significance of Lipids to Biofilm Formation in Candida albicans: An Emerging Perspective. J. Fungi 4 (2018).

48. L. Scorzoni et al., Antifungal Therapy: New Advances in the Understanding and Treatment of Mycosis. Front. Microbiol. 8 (2017).

49. M. J. Hynes, S. L. Murray, A. Andrianopoulos, M. A. Davis, Role of Carnitine Acetyltransferases in Acetyl Coenzyme A Metabolism in Aspergillus nidulans. Eukaryot. Cell 10, 547–555 (2011).

50. M. C. Lorenz, Carbon Catabolite Control in Candida albicans: New Wrinkles in Metabolism. mBio 4, e00034–00013 (2013).

51. S. Sánchez et al., Carbon source regulation of antibiotic production. J. Antibiot. 63, 442–459 (2010).

52. H. Yang et al., Sterol Esterification in Yeast: A Two-Gene Process. Science 272, 1353–1356 (1996).

53. Y. Wang et al., Antimicrobial Blue Light Inactivation of Gram-Negative Pathogens in Biofilms: In Vitro and In Vivo Studies. J. Infect. Dis. 213, 1380–1387 (2016).

54. P. G. Pappas, M. S. Lionakis, M. C. Arendrup, L. Ostrosky-Zeichner, B. J. Kullberg, Invasive candidiasis. Nature Reviews Disease Primers 4, 18026 (2018).

55. C. A. Kumamoto, Candida biofilms. Curr. Opin. Microbiol. 5, 608–611 (2002).

56. G. Ramage, E. Mowat, B. Jones, C. Williams, J. Lopez-Ribot, Our Current Understanding of Fungal Biofilms. Crit. Rev. Microbiol. 35, 340–355 (2009).

57. S. Silva, C. F. Rodrigues, D. Araújo, M. E. Rodrigues, M. Henriques, Candida Species Biofilms’ Antifungal Resistance. J Fungi (Basel) 3, 8 (2017).

58. J. E. Nett, Future directions for anti-biofilm therapeutics targeting Candida. Expert Review of Anti-infective Therapy 12, 375–382 (2014).

59. A. A. Lattif et al., Lipidomics of Candida albicans biofilms reveals phase-dependent production of phospholipid molecular classes and role for lipid rafts in biofilm formation. Microbiology 157, 3232–3242 (2011).

60. D. Alim, S. Sircaik, S. L. Panwar, The Significance of Lipids to Biofilm Formation in Candida albicans: An Emerging Perspective. J. Fungi 4, 140 (2018).

61. A. Singh, R. Prasad, Comparative Lipidomics of Azole Sensitive and Resistant Clinical Isolates of Candida albicans Reveals Unexpected Diversity in Molecular Lipid Imprints. PLOS ONE 6, e19266 (2011).

62. A. Singh, V. Yadav, R. Prasad, Comparative Lipidomics in Clinical Isolates of Candida albicans Reveal Crosstalk between Mitochondria, Cell Wall Integrity and Azole Resistance. PLOS ONE 7, e39812 (2012).

63. J. Gao et al., Candida albicans gains azole resistance by altering sphingolipid composition. Nat. Commun. 9, 4495 (2018).

64. M. Connerth et al., Oleate Inhibits Steryl Ester Synthesis and Causes Liposensitivity in Yeast*. J. Biol. Chem. 285, 26832–26841 (2010).

65. D. Fu, G. Holtom, C. Freudiger, X. Zhang, X. S. Xie, Hyperspectral Imaging with Stimulated Raman Scattering by Chirped Femtosecond Lasers. J. Phys. Chem. B 117, 4634–4640 (2013).

66. B. Liu et al., Label-free spectroscopic detection of membrane potential using stimulated Raman scattering. Appl. Phys. Lett. 106, 173704 (2015).

67. J. Folch, M. Lees, G. H. S. Stanley, A Simple method for the isolation and purification of total lipides from animal tissues. J. Biol. Chem. 226, 497–509 (1957).

68. G. Liebisch et al., High throughput quantification of cholesterol and cholesteryl ester by electrospray ionization tandem mass spectrometry (ESI-MS/MS). Biochim. Biophys. Acta 1761, 121–128 (2006).

