## Supporting Information for "Fingerprint SRS Imaging Unveils Ergosteryl Ester as a Metabolic Signature of Azole-Resistant *Candida albicans*"

**This PDF file includes:**

Figures S1 to S12
Tables S1
SI References

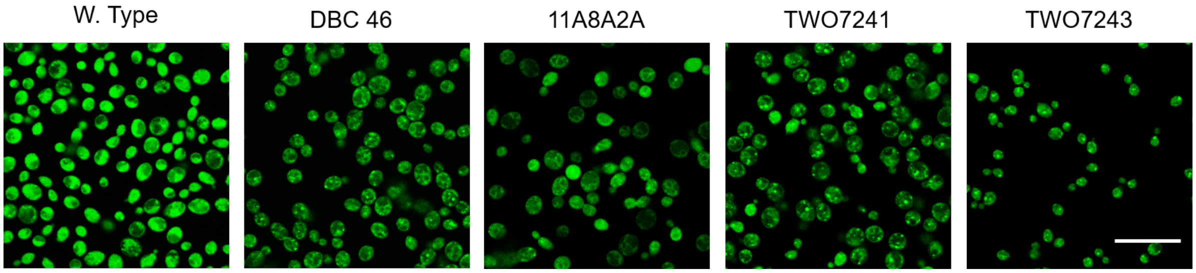


**Fig. S1.** Confocal fluorescence imaging of lipids stained by BODIPY dye in *C. albicans* W. Type, DBC 46, 11A8A2A, TWO7241, and TWO7243. Bar scale represents 10 μm.

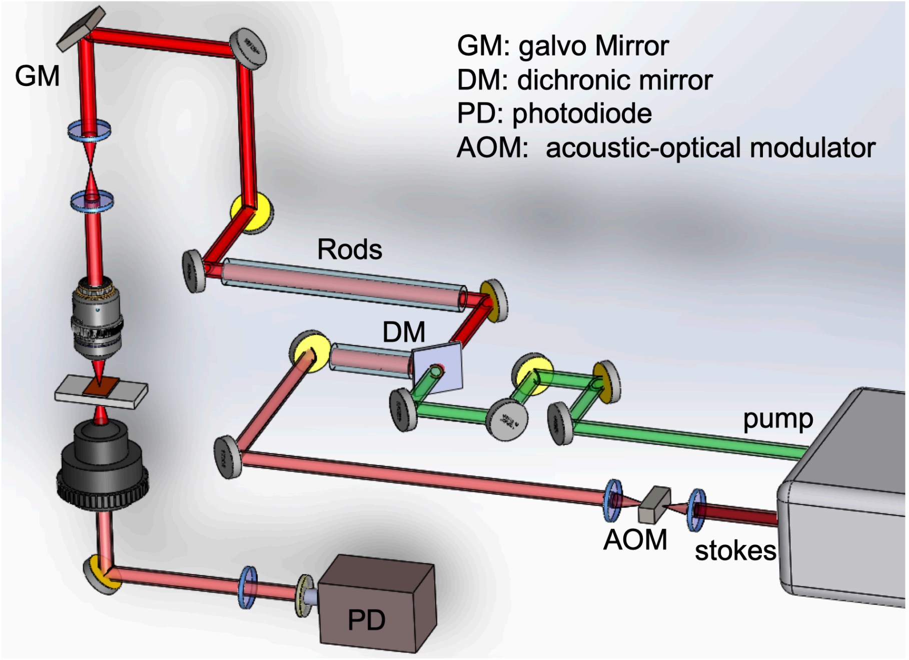


**Fig. S2.** Hyperspectral SRS imaging setup. A femtosecond laser provided the pump and Stokes laser source. The Stokes beam was modulated by an acousto-optic modulator (AOM). After combination through a dichroic mirror (DM), both the pump and Stokes beams were chirped by glass rods and then sent to a laser-scanning microscope equipped with galvo mirror (GM). The signal was collected by an oil condenser and was sent to a photodiode (PD).

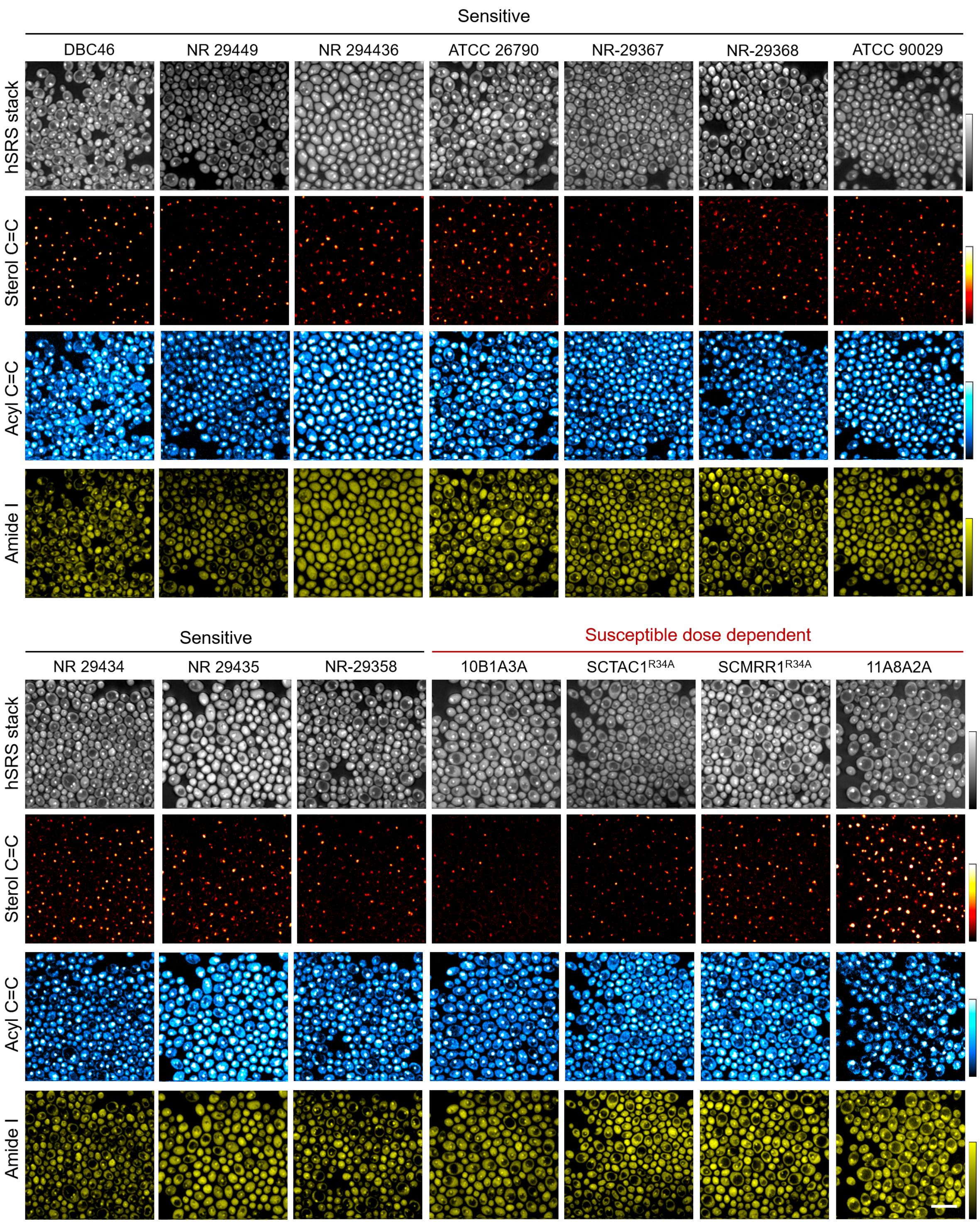


**Fig. S3**. hSRS imaging of multiple fluconazole-susceptible and susceptible dose dependent *C. albicans*. Bar scale represents 10 μm.

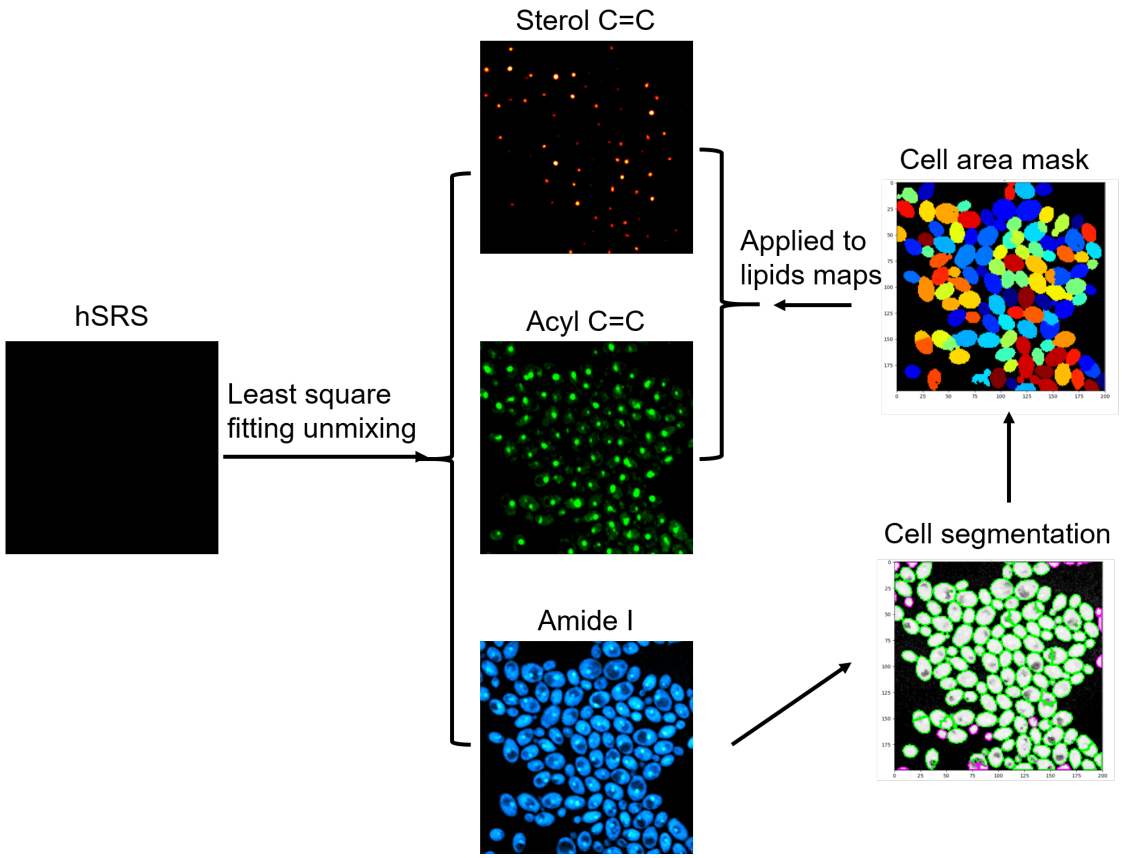


**Fig. S4**. Segmentation of the decomposed concentration maps of the hyperspectral SRS images to generate maps of intracellular proteins, quantification of sterol C=C and acyl C=C levels based on the signal threshold adjustment.

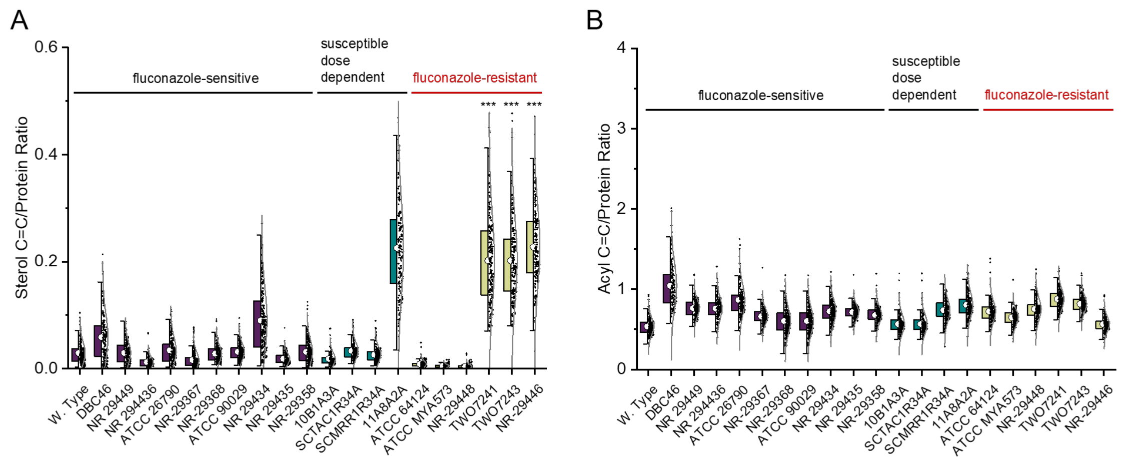


**Fig. S5.** EE to protein ratio intensity (A) and acyl C=C to protein ratio intensity (B). Significance was evaluated using an unpaired *t*-test (***, $p < 0.001$).

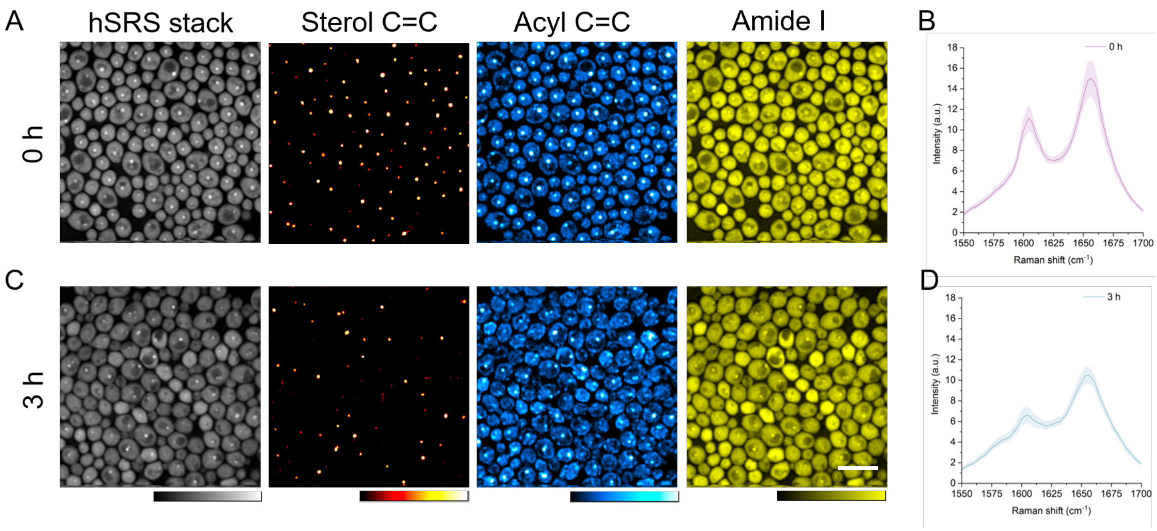


**Fig. S6**. Hypespectral SRS imaging unmixing showing decreased EE accumulation in stationary phase azole-resistant *C. albicans* TWO7241 (A) before and (C) after incubation in fresh YPD medium for 3 h. hSRS spectra of lipids accumulated in *C. albicans* TWO7241 (B) before and (D) after incubation in fresh medium for 3 h. Bar scale represents 10 μm.

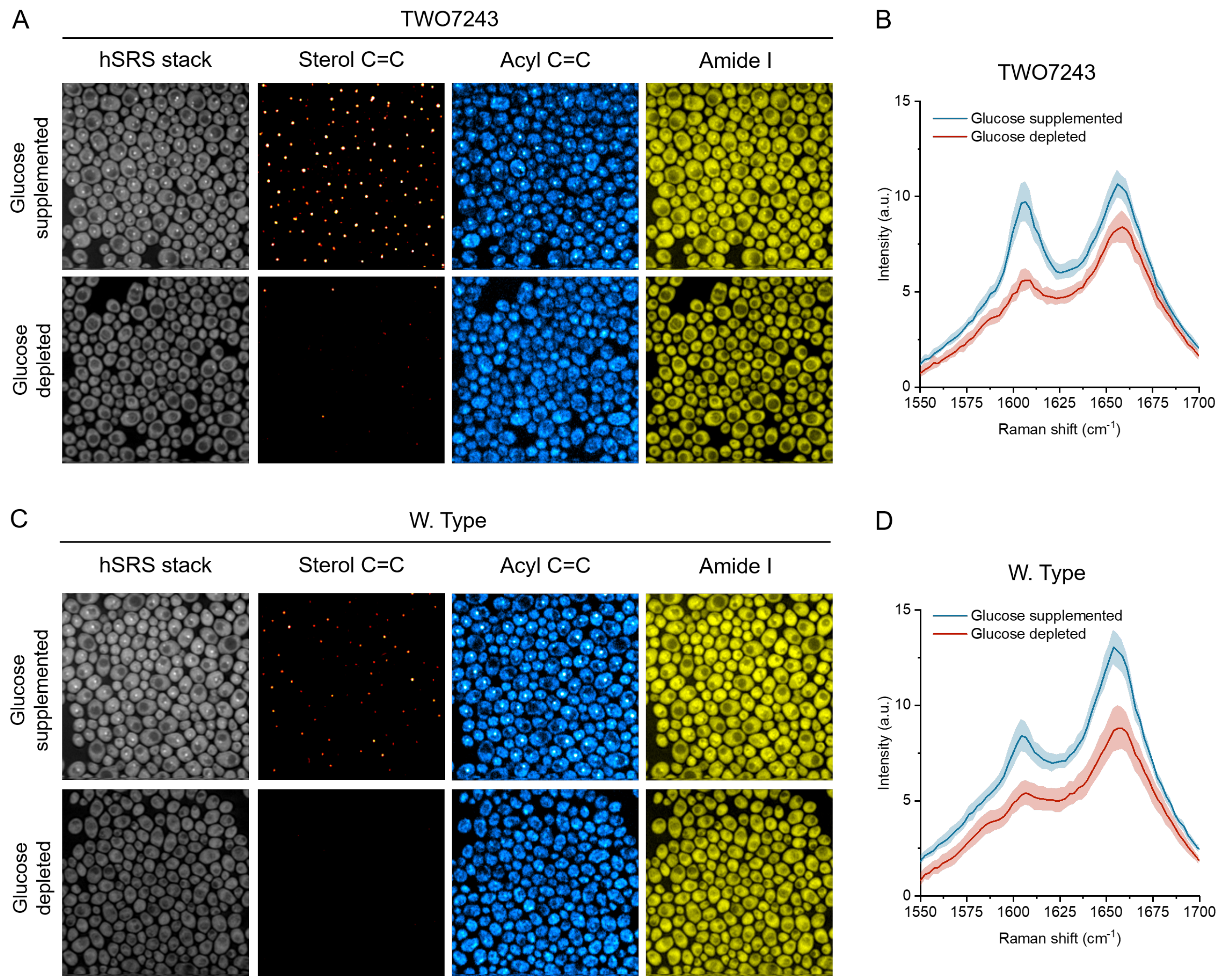


**Fig. S7.** LDs intensity level is reduced in glucose supplemented medium compared to glucose depleted medium in azole-resistant *C. albicans* TWO7243 (A, B) azole-susceptible *C. albicans* W. Type (C, D).

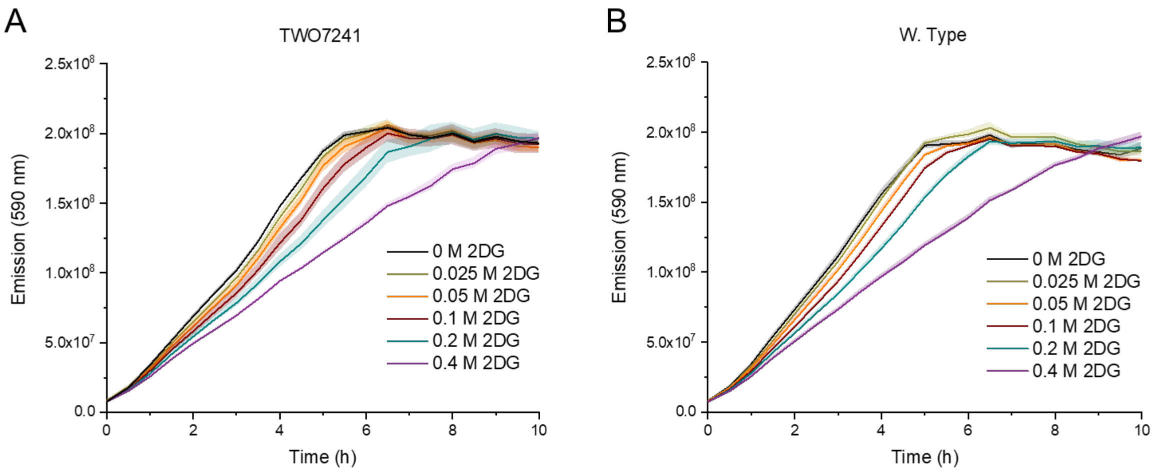


**Fig. S8.** 2DG toxicity study on azole-resistant *C. albicans* TWO7241 (A) and azole-susceptible *C. albicans* W. Type (B).

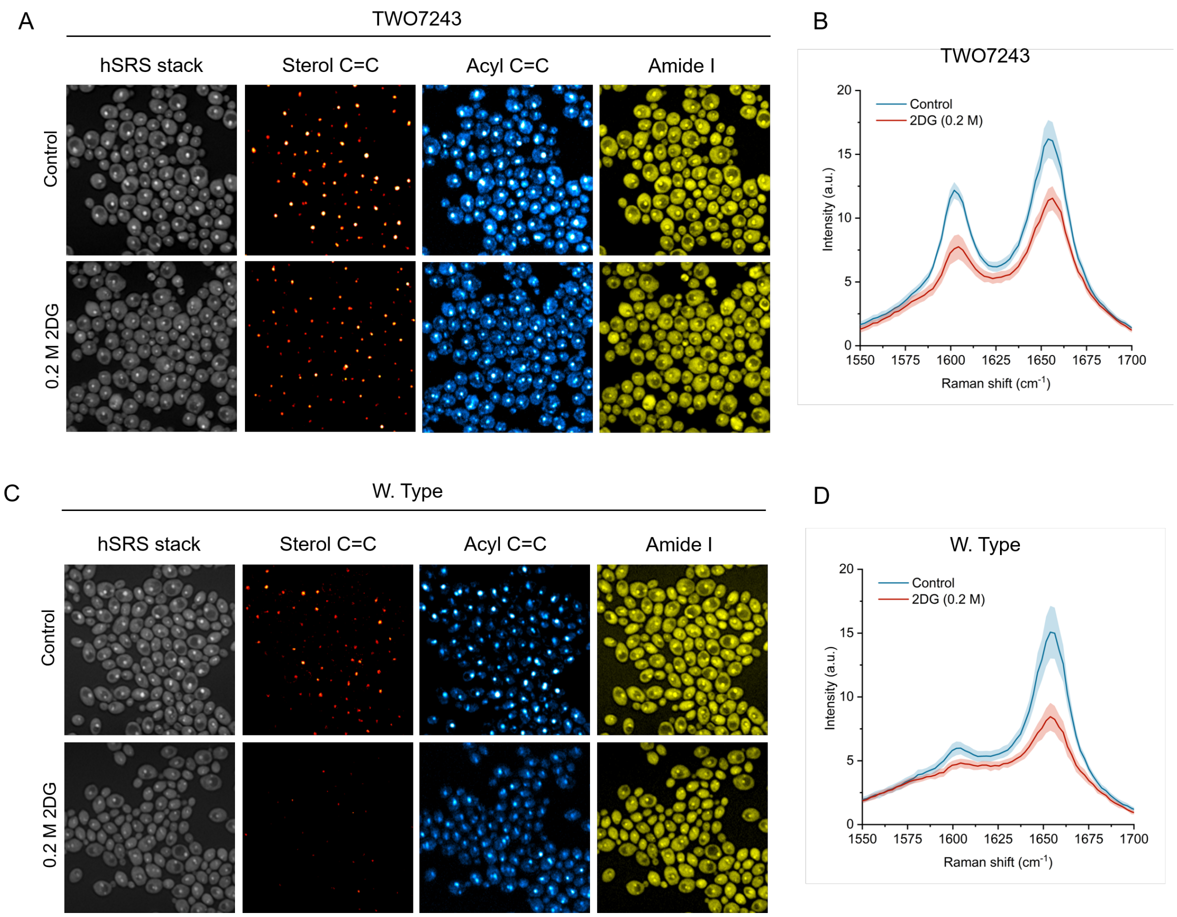


**Fig. S9.** LDs intensity level is reduced under 2DG treatment compared to non-treated group in azole-resistant *C. albicans* TWO7243 (A, B) and azole-susceptible *C. albicans* W. Type (C, D).

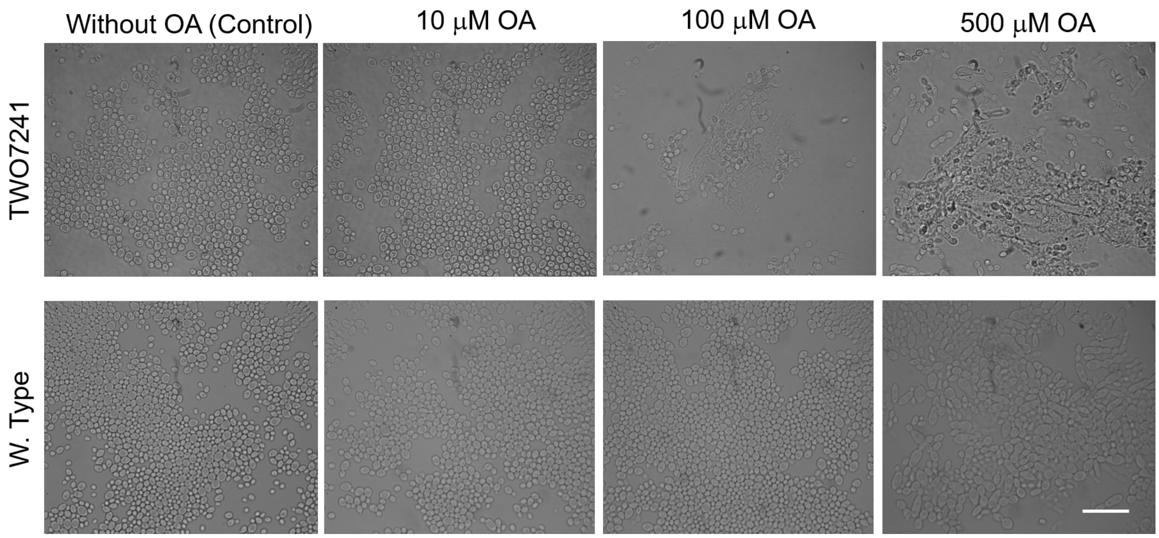


**Fig. S10.** Transmission images of concentration-varied OA treated azole-resistant *C. albicans* TWO7241 and azole-sensitive *C. albicans* W. Type. Bar scale represents 10 μm.

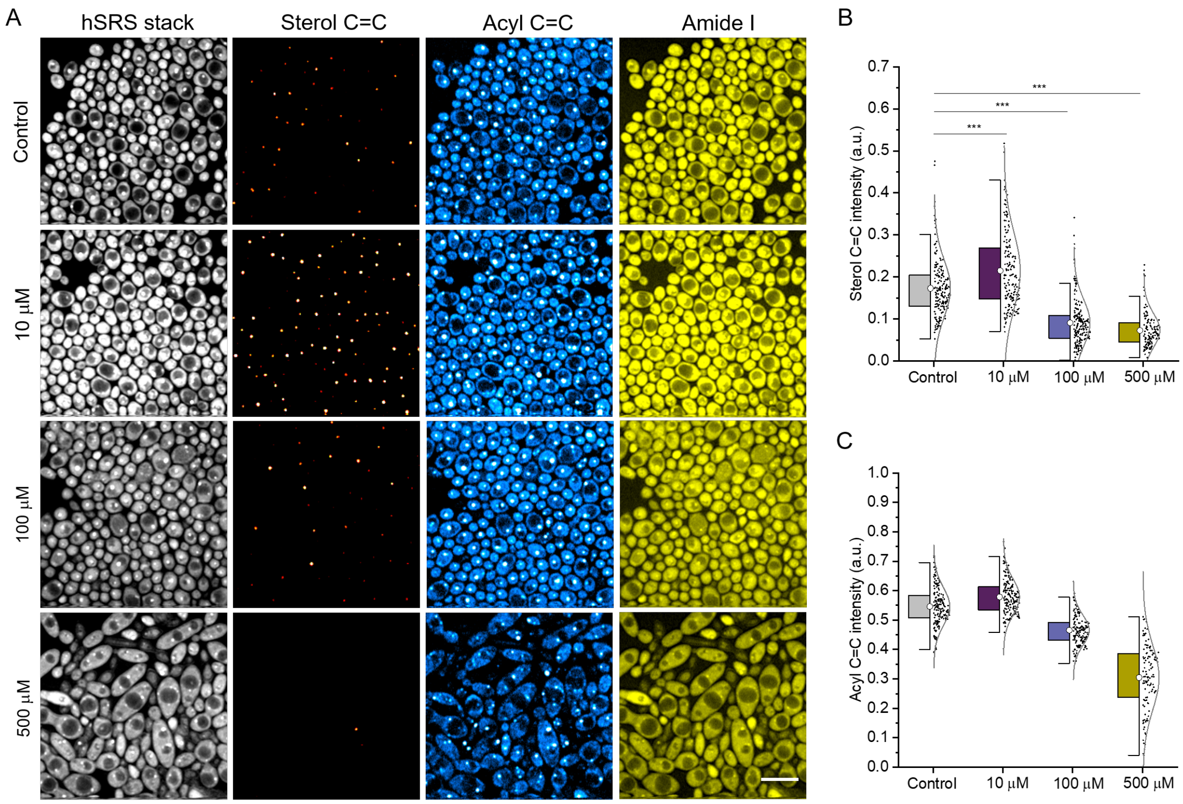


**Fig. S11.** EE accumulation in azole-sensitive *C. albicans* W. Type is less sensitive to oleate. (A) Fingerprinting hyperspectral SRS images of *C. albicans* W. Type under concentration-dependent oleate treatment. Quantification of (B) EE and (C) acyl C=C levels in hSRS unmixed concentration maps. Significance was measured using unpaired t-test (*, $p < 0.05$; **, $p < 0.01$; ***, $p < 0.001$). Bar scale represents 10 μm.

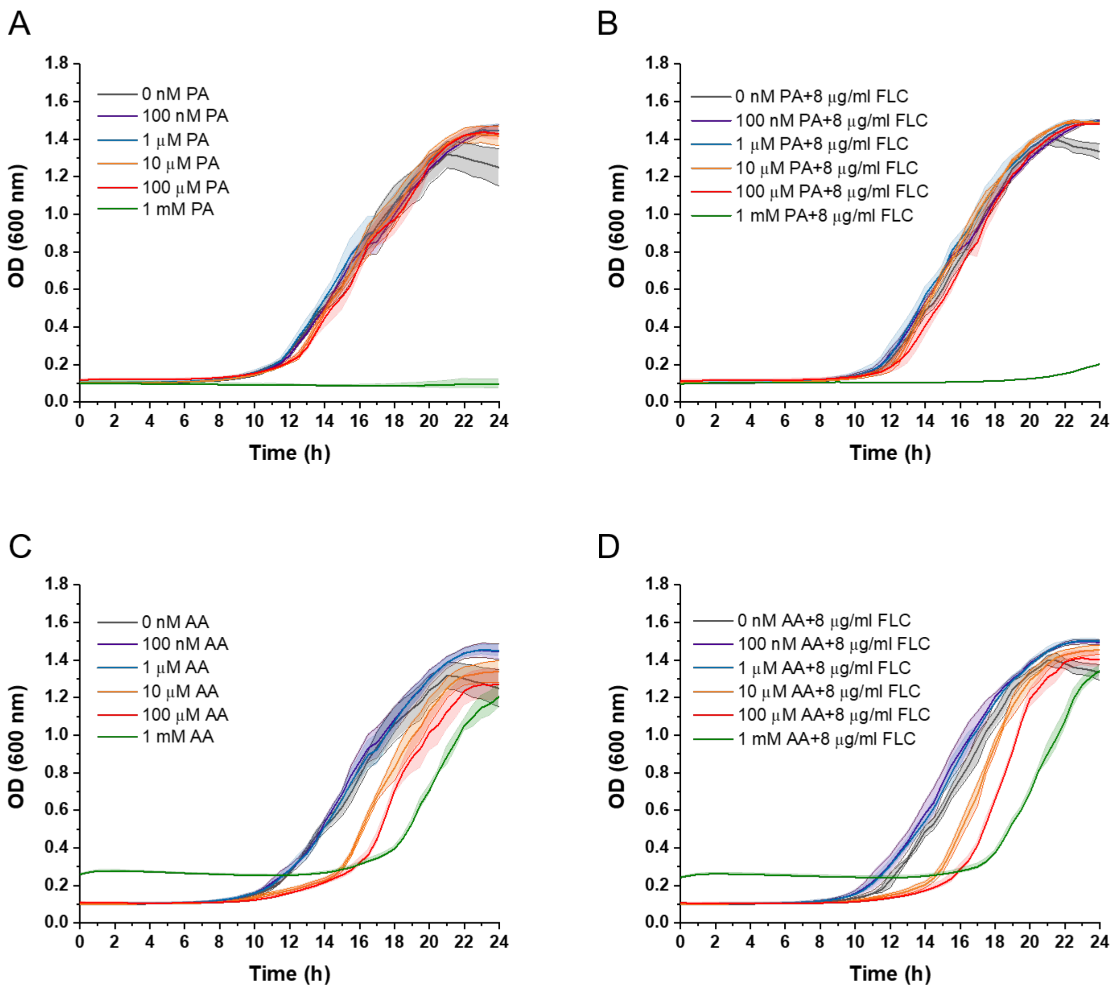


**Fig. S12.** PA (A, B) and AA (C, D) do not show synergistic effect with fluconazole on inhibition cell growth of *C. albicans* TWO7241.

**Table S1**. **List of *C. albicans* strains used in this study and the MICs of fluconazole against these strains.**

| *C. albicans* Strain ID | Fluconazole MIC (µg/ml) | Description | Mechanism of azole resistance | References |
|---|---|---|---|---|
| SC5314 | 0.125 | Fluconazole-sensitive mutant strain | Not applicable | (1) |
| DBC 46 | 0.0625 | Fluconazole-sensitive mutant strain | Reduced ergosterol production, a heterozygous deletion in *ERG11* | |
| 11A8A2A | 2 | Susceptible dose dependent strain | Homozygous activating mutation in *UPC2* (*ERG11* overexpression) | (2) |
| TWO7243 | 64 | Fluconazole resistant strain | Overexpression of *CDR1* and *ERG11* | (3) |
| TWO7241 | 32 | Fluconazole resistant strain | Overexpression of *MDR1* and *ERG11* | (3) |
| NR 29434 | 0.125 | Fluconazole-sensitive strain | Not applicable | (4) |
| NR 29449 | 0.125 | Fluconazole-sensitive strain | Not applicable | (4) |
| NR 294436 | 0.125 | Fluconazole-sensitive strain | Not applicable | (5) |
| NR 29435 | 0.125 | Fluconazole-sensitive strain | Not applicable | (4) |
| ATCC 90029 | 0.125 | Fluconazole-sensitive strain | Not applicable | (6) |
| ATCC 26790 | 0.5 | Fluconazole-sensitive strain showing a significant trailing growth | Not applicable | (4) |
| NR 29358 | 0.5 | Fluconazole-sensitive strain showing a significant trailing growth | Not applicable | (6) |
| NR 29367 | 0.5 | Fluconazole-sensitive strain showing a significant trailing growth | Not applicable | (6) |
| NR 29368 | 0.5 | Fluconazole-sensitive strain showing a significant trailing growth | Not applicable | (6) |
| 10B1A3A | 4 | Susceptible dose dependent strain | Homozygous single mutation in *ERG11* ($ERG11^{K143R}$) | (7) |
| $SCTAC1^{R34A}$ | 4 | Susceptible dose dependent strain | Homozygous activating mutation in *TAC1* (*CDR1* and *CDR2* overexpression) | (8) |
| $SCMRR1^{R34A}$ | 4 | Susceptible dose dependent strain | Homozygous activating mutation in *MRR1* (*MDR1* overexpression) | (9) |

| | | | | |
|---|---|---|---|---|
| NR 29446 | 64 | Fluconazole resistant strain | Unknown | (10) |
| NR 29448 | > 128 | Fluconazole resistant strain | Unknown | (6) |
| ATCC MYA573 | 128 | Fluconazole resistant strain | Unknown | (11, 12) |
| ATCC 64124 | 64 | Fluconazole resistant strain | Multiple mutations in *ERG11* | (6, 12) |

### SI References


1. H. E. Eldesouky *et al.*, Repurposing approach identifies pitavastatin as a potent azole chemosensitizing agent effective against azole-resistant *Candida* species. *Sci. Rep.* **10**, 7525 (2020).
2. S. A. Flowers *et al.*, Gain-of-function mutations in UPC2 are a frequent cause of ERG11 upregulation in azole-resistant clinical isolates of *Candida albicans*. *Eukaryot. Cell* **11**, 1289-1299 (2012).
3. K. Koselny *et al.*, The Celecoxib Derivative AR-12 Has Broad-Spectrum Antifungal Activity In Vitro and Improves the Activity of Fluconazole in a Murine Model of Cryptococcosis. *Antimicrob Agents Chemother* **60**, 7115-7127 (2016).
4. S. Thangamani *et al.*, Ebselen exerts antifungal activity by regulating glutathione (GSH) and reactive oxygen species (ROS) production in fungal cells. *Biochim. Biophys. Acta - Gen. Subj.* **1861**, 3002-3010 (2017).
5. A. AbdelKhalek, C. R. Ashby, Jr., B. A. Patel, T. T. Talele, M. N. Seleem, In Vitro Antibacterial Activity of Rhodanine Derivatives against Pathogenic Clinical Isolates. *PLoS One* **11**, e0164227 (2016).
6. H. E. Eldesouky, A. Mayhoub, T. R. Hazbun, M. N. Seleem, Reversal of Azole Resistance in *Candida albicans* by Sulfa Antibacterial Drugs. *Antimicrob Agents Chemother* **62** (2018).
7. A. T. Nishimoto *et al.*, In Vitro Activities of the Novel Investigational Tetrazoles VT-1161 and VT-1598 Compared to the Triazole Antifungals against Azole-Resistant Strains and Clinical Isolates of *Candida albicans*. *Antimicrob Agents Chemother* **63** (2019).
8. B. Ramírez-Zavala, H. Manz, F. Englert, P. D. Rogers, J. Morschhäuser, A Hyperactive Form of the Zinc Cluster Transcription Factor Stb5 Causes YOR1 Overexpression and Beauvericin Resistance in *Candida albicans*. *Antimicrob Agents Chemother* **62** (2018).
9. S. Schubert *et al.*, Regulation of efflux pump expression and drug resistance by the transcription factors Mrr1, Upc2, and Cap1 in *Candida albicans*. *Antimicrob Agents Chemother* **55**, 2212-2223 (2011).
10. H. Mohammad *et al.*, Discovery of a Novel Dibromoquinoline Compound Exhibiting Potent Antifungal and Antivirulence Activity That Targets Metal Ion Homeostasis. *ACS Infect Dis* **4**, 403-414 (2018).
11. H. E. Eldesouky *et al.*, Repurposing approach identifies pitavastatin as a potent azole chemosensitizing agent effective against azole-resistant *Candida* species. *Sci Rep* **10**, 7525 (2020).
12. H. Mohammad, H. E. Eldesouky, T. Hazbun, A. S. Mayhoub, M. N. Seleem, Identification of a Phenylthiazole Small Molecule with Dual Antifungal and Antibiofilm Activity Against *Candida albicans* and *Candida auris*. *Sci Rep* **9**, 18941 (2019).
